# E- and N-cadherin drive hepatic polarity and lumen elongation via opposing effects on RhoA activity

**DOI:** 10.1101/2025.10.06.680681

**Authors:** Junya Hayase, Li Yang, Yu-Heng Zhou, Kangji Wang, Cheng-Ran Xu, Erfei Bi

## Abstract

Hepatocytes exhibit distinct polarity, forming narrow apical tubes (bile canaliculi, BCs) between adjacent cells. These structures essential to liver architecture and function. Unlike most epithelial cells, hepatocytes express both E- and N-cadherin but their functions and mechanisms remain unknown. We show that E- and N-cadherin are collectively required for hepatic polarity and BC formation but act through distinct, spatially segregated pathways. Both localize to adherens junctions; E-cadherin additionally localizes to lateral membranes and the cleavage furrow during cell division, where it promotes BC elongation and new cell-cell contact formation by controlling spindle orientation and RhoA activation via NuMA and ARHGEF17. N-cadherin maintains hepatic polarity by facilitating RhoA inactivation through the p120-catenin family member ARVCF and its partner p190B/ARHGAP5. Thus, dual cadherin expression drives hepatic polarity and BC formation by controlling RhoA activity in a coordinated but opposing manner.

**Summary:** Hayase et al. show that E-cadherin promotes bile canaliculi elongation via RhoA activation and oriented cell division, while N-cadherin maintains hepatic polarity by suppressing RhoA, revealing the function and mechanism of dual cadherin expression in hepatocytes.

## Introduction

The function of many internal organs such as the kidney, small intestine, and the liver depends on the generation of an epithelium (Buckley and St Johnston, 2022; Garcia et al., 2018). The epithelial tissue consists of epithelial cells that are tightly interconnected through several types of junctional complexes including adherens junctions (AJs), tight junctions (TJs), and desmosomes, which enable the tissue to form sheet-like structures (Garcia et al., 2018). Among these junctions, AJs play a principal role in cell-cell adhesion and in the rearrangement and movement of epithelial tissues (Campas et al., 2024; Guillot and Lecuit, 2013; Takeichi, 2014). The epithelial cadherin (E-cadherin) is a key component for the AJ (Takeichi, 2014), and links cell-cell contacts to circumferential actomyosin networks underneath the apical membrane, through its binding partners catenins, thus contributing to the integrity of epithelial tissues (Campas et al., 2024; Guillot and Lecuit, 2013).

The stereotyped architecture of epithelial cell sheets, however, is disrupted during developmental or pathological processes including Epithelial to Mesenchymal Transition (EMT) (Peglion and Etienne-Manneville, 2024; Yang et al., 2020). In cells undergoing EMT, E-cadherin is downregulated while neuronal cadherin (N-cadherin), another type of classical cadherin expressed in non-epithelial cells, is often upregulated (Loh et al., 2019; van Roy, 2014). Although both E-cadherin and N-cadherin are engaged in cell-cell adhesion, the switch from E- to N-cadherin is associated with changes in cell behavior including motility (Loh et al., 2019) and in cell polarity (Scarpa et al., 2015). Despite extensive studies of these cadherins in different settings including EMT and tumor metastasis (Loh et al., 2019; Radice, 2013; van Roy, 2014), the molecular mechanisms underlying their distinct roles are still lacking.

The liver is the largest internal organ responsible for vital functions including metabolism, detoxification, blood glucose homeostasis, serum protein synthesis, and bile production (Lotto et al., 2023). These functions of the liver rely on its elaborate architecture formed during development (Lotto et al., 2023; Schulze et al., 2019). Hepatocytes are the parenchymal cells of epithelial origin in the liver and are organized into cords in which two adjacent cells share a narrow apical lumen known as the bile canaliculus (BC). BC serves as the site for bile excretion (Schulze et al., 2019; Treyer and Musch, 2013). This apical lumen is continuous with adjoining hepatocytes and extends throughout the hepatic cords, forming a network of bile canaliculi that eventually connect to the bile ducts (Tanimizu and Mitaka, 2017; Treyer and Musch, 2013). The basal surfaces of hepatocytes have the unique feature of being flanked by sinusoidal spaces that contain only sparse connective tissue chiefly consisting of collagen fibers, allowing hepatocytes access to sinusoidal blood flow, which is in marked contrast to the basal membranes of most epithelial cell types that contact a dense basement membrane composed primarily of laminin and collagen IV (Tanimizu and Mitaka, 2017; Treyer and Musch, 2013). Thus, the elaborate architecture and vital function of the liver are highly dependent on the unique cell polarity of hepatocytes. However, the mechanisms that establish and maintain this polarity, particularly during BC formation and elongation, remain unclear.

Unlike other epithelial cells, hepatocytes express both E- and N-cadherin (Doi et al., 2007; Straub et al., 2011), and their expression is developmentally regulated (Doi et al., 2007). In mouse liver, both cadherins are present in all hepatoblasts/hepatocytes during embryogenesis, but postnatally their distribution becomes zonally restricted: E-cadherin is enriched in the peri- portal vein region with a sharp boundary, where N-cadherin is expressed throughout the liver lobule but concentrated in the peri-central vein region (Doi et al., 2007). During liver regeneration, hepatocytes that normally lack E-cadherin are induced to express it, resulting in dual cadherin expression (He et al., 2021; Lin et al., 2023; Wang et al., 2024). Thus, co-expression of E- and N-cadherin is a hallmark of both developing and regenerating hepatocytes. However, whether these cadherins have distinct or overlapping roles in hepatic polarity and BC development remains unknown.

In this study, we use the rat hepatocyte cell line Can 10 (Peng et al., 2006; Treyer and Musch, 2013; Wang et al., 2014), which resembles developing hepatocytes, to show that although E- and N-cadherin are collectively required for hepatic polarity and BC formation, they play distinct roles: E-cadherin promotes BC elongation by coordinating mitotic spindle orientation and spatiotemporal activation of RhoA through NuMA (Kiyomitsu and Boerner, 2021) and ARHGEF17. N-cadherin maintains hepatic polarity by suppressing RhoA activity via the p120-catenin family member ARVCF (Mariner et al., 2000; Sirotkin et al., 1997) and its partner ARHGAP5/p190B (Cho et al., 2010). Thus, both cadherins act independently yet cooperatively to ensure hepatic polarity and tubular BC formation—processes essential for liver architecture and function.

## Results

### A proper ratio of E- and N-cadherin in hepatocytes is critical for hepatic polarity development

To investigate the role of cadherins in AJ assembly and hepatic polarity development, we examined their expression and localization during mouse liver development. Consistent with a previous report (Doi et al., 2007), we found that both E-cadherin and N-cadherin are expressed in hepatoblasts and hepatocytes from embryonic day 13.5 (E13.5) to postnatal day (P0), corresponding to the stage of bile canaliculi (BC) formation, elongation, and branching (Fig. 1, A and B; and Fig. S1 A). Each hepatocyte expresses both cadherins, which delineate BC structures enclosed by ZO-1 (Fig. 1 A). In contrast, E- and N-cadherin exhibited zonal distribution after the BC network was established (Fig. S1 B) (Doi et al., 2007).

**Figure 1.**
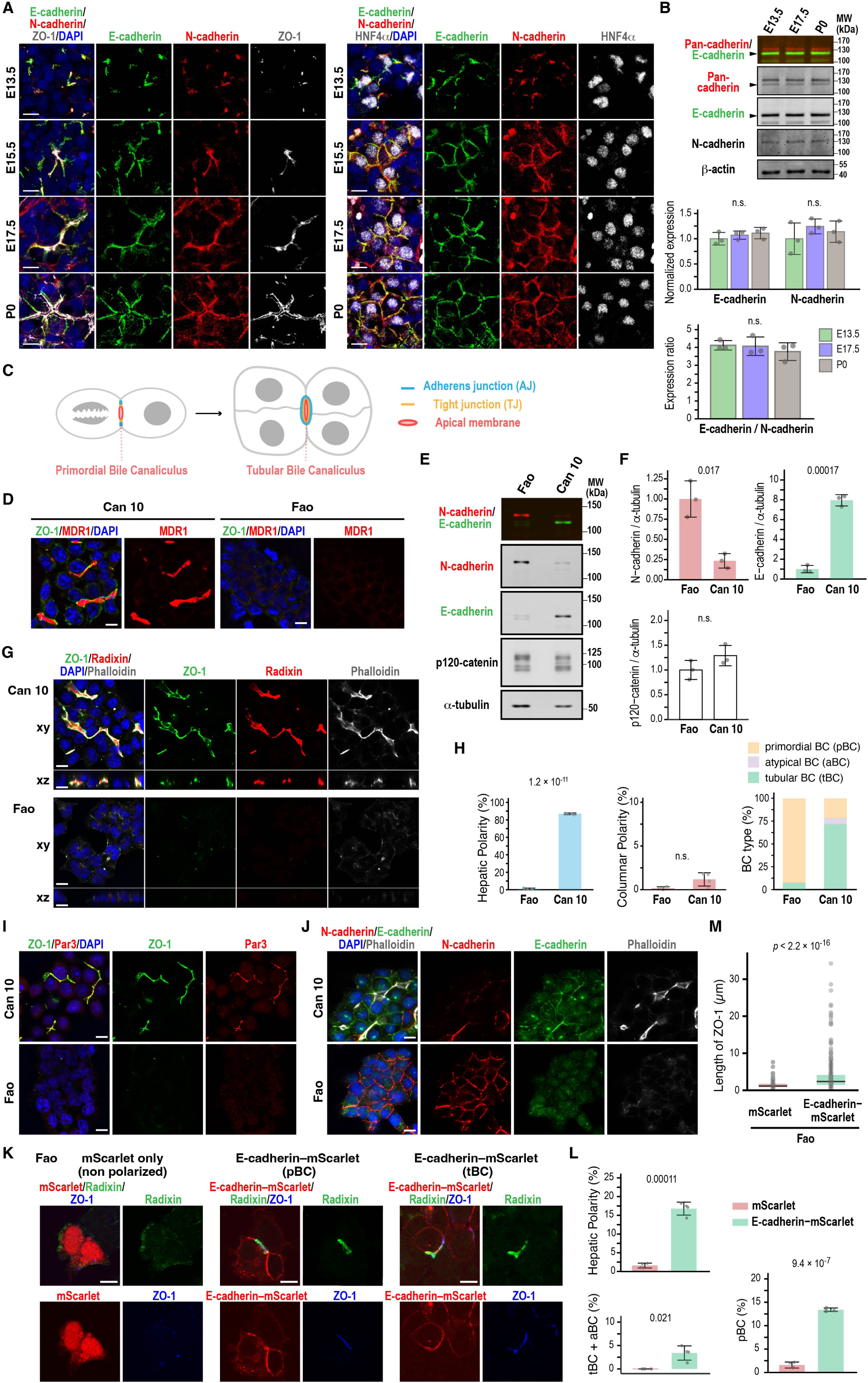
Increased E-cadherin expression relative to N-cadherin induces hepatic polarity. (A) Expression of E- and N-cadherin in hepatoblasts and hepatocytes during mouse liver development. Liver sections from different developmental stages were immunostained with antibodies against E-cadherin, N-cadherin, ZO-1, and HNF4α along with DAPI staining. (B) Immunoblot analysis of hepatoblasts and hepatocytes at different developmental stages of the mouse liver. Immunoblotting was performed using antibodies against E-, N-, and pan- cadherin, as well as β-actin. Arrowheads indicate E-cadherin corresponding to the lower band of pan-cadherin. Molecular weights (MWs) of marker proteins are indicated in kDa. Middle: Normalized expression of E- and N-cadherin during BC biogenesis; the values at E13.5 are set to 1.0. Bottom: Expression ratio of E-cadherin to N-cadherin during BC biogenesis after normalizing pan-cadherin antibody affinities to E- and N-cadherin. Data represent means ± SD from three independent samples. See also Figure S1, C and D. (C) Schematic of tubular BC formation. Oriented cell division of a cell with a primordial BC partitions the pre-existing lumen into two daughter cells, resulting in a tubular BC surrounded by three or more cells. (D) Representative instant Structured Illumination Microscopy (iSIM) images of polarized Can 10 cells and unpolarized parental Fao cells. Cells were cultured for 3 days, then fixed and stained with DAPI and antibodies against MDR1 and ZO-1. (E) Immunoblot analysis of E- and N-cadherin expression in Fao and Can 10 cells. Immunoblotting was performed using antibodies against N-cadherin, E-cadherin, p120-catenin, and α-tubulin. Molecular weights (MWs) of marker proteins are indicated in kDa. (F) Normalized intensity ratios of N-cadherin, E-cadherin, and p120-catenin relative to tubulin in Fao and Can 10 cells. Values represent means ± SD from three independent experiments shown in (E). Ratios in Fao cells were set to 1.0. See also Figure S1, D and E. (G) Analysis of cell polarization in Can 10 and Fao cells. Cells were cultured for 3 days, then fixed and stained with phalloidin, DAPI, and antibodies against radixin and ZO-1. iSIM images were shown in horizontal (xy) and orthogonal (xz) views. (H) Quantification of Can 10 and Fao cells with various types of polarity and BC structures. Values are means ± SD from four independent experiments (≥401 cells counted per sample). (I) Analysis of TJ protein localization and assembly in Can 10 and Fao cells. Cells were cultured for 3 days, then fixed and stained with DAPI and antibodies against Par-3 and ZO-1. Representative iSIM images are shown. (J) Analysis of AJ protein localization and assembly in relation to F-actin in Can 10 and Fao cells. Cells were cultured for 3 days, then fixed and stained with phalloidin, DAPI, and antibodies against E-cadherin and N-cadherin. Representative iSIM images are shown. (K) Induction of hepatic polarity in Fao cells by increasing E-cadherin expression. Fao cells were cultured for 2 days following transduction with lentiviruses expressing either mScarlet or E-cadherin–mScarlet, then fixed and stained with antibodies against radixin and ZO-1. Representative confocal images of unpolarized cells expressing mScarlet only, or polarized cells with primordial BC (pBC) or tubular BC (tBC) expressing E-cadherin–mScarlet, are shown. (L) Quantification of Fao cells expressing mScarlet alone or E-cadherin–mScarlet with indicated polarity and BC structures. Values represent means ± S.D. from four independent experiments (≥154 cells counted per sample). (M) Quantification of ZO-1 length in Fao cells expressing mScarlet only or E-cadherin– mScarlet. Values are from four independent experiments (≥211 ZO-1-positive structures counted per condition). Scale bars, 10 µm (A, D, G, and I–K). p-values are indicated at the top of each graph. n.s., not significant.

Western blotting with E- and N-cadherin-specific and pan-cadherin antibodies showed that hepatoblasts and hepatocytes isolated from E13.5, E17.5, and P0 expressed both cadherins, and their levels and ratios did not vary significantly during these developmental stages (Fig. 1 B). These findings raise the question of whether co-expression of E- and N-cadherin is critical for establishing and/or maintaining the unique hepatic polarity. To address this, we used the rat hepatocyte cell line Can 10, which recapitulates the BC developmental stage observed *in vivo* (Fig. 1, C and D) (Wang et al., 2014). We first compared the protein levels of these cadherins between Can 10 cells and their parental Fao cells, which lack hepatic polarity (Peng et al., 2006). Similar to hepatocytes in developing livers (Fig. 1, A and B), both cadherins were expressed in Can 10 and Fao cells (Fig. 1, E and F). However, E-cadherin was predominant in Can 10, while N-cadherin was predominant in Fao (Fig. 1, E and F). Using GFP-tagged E- and N-cadherin to normalize pan-cadherin antibody affinities, we found that the E-to-N-cadherin expression ratio in Can 10 cells (∼4.6) closely matched that in hepatoblasts/hepatocytes (∼4.0) from E13.5 to P0, favoring E-cadherin expression (Fig. 1 B; Fig. S1, C and D). The total level of p120-catenin—a binding partner of both E- and N-cadherin—was comparable in Can 10 and Fao, with only a difference in the expression of its mesenchymal splice variant, p120-1 (Fig. 1, E and F; Fig. S1 E) (Keirsebilck et al., 1998; Mo and Reynolds, 1996). Approximately 80% of Can 10 cells exhibited hepatic polarity, defined as a radixin-positive apical domain enclosed by two or more cells (Fig. 1, G and H). More than half of these polarized cells formed tubular BCs involving three cells or more cells—structures similar to those observed in hepatocytes at E 15.5 and E17.5 (Fig. 1, A and G). In contrast, Fao cells lacked both hepatic polarity and typical epithelial columnar polarity, characterized by apical membranes facing to the free surface (Fig. 1, G–J). This was despite their abundant N-cadherin expression and apparent cadherin-mediated cell-cell contacts (Fig. 1 J). The high E/N-cadherin ratio and strong polarization capacity of Can 10 cells prompted us to test whether forced E-cadherin expression could induce polarity in Fao cells. Strikingly, E-cadherin– mScarlet expression induced hepatic polarity (Fig. 1, K and L), elongating both ZO-1-positive junctions (Fig. 1 M) and BC structures (Fig. 1, K and L). Thus, in the presence of N-cadherin, upregulating E-cadherin is sufficient to induce hepatic polarity and drive BC formation and elongation. Together, these observations suggest that a proper ratio of E- and N-cadherin in hepatocytes is critical for establishing hepatic polarity.

### Distinct and shared roles of E- and N-cadherin in hepatic polarity development and BC biogenesis

To define their specific roles in hepatocytes, we depleted E- and N-cadherin individually or together in Can 10 cells using siRNAs. The transient depletion of E-cadherin markedly reduced the formation of tubular BCs (Fig. 2, A–C). In contrast, cells exhibiting hepatic polarity or apical-basal polarity— which includes both hepatic and columnar polarity—showed only a modest but significant reduction (Fig. 2 C; and Fig. S2 A). Most E-cadherin-knockdown cells formed only primordial BCs between two opposing cells (Fig. 2, B and C), suggesting that E-cadherin is primarily required for BC elongation. N-cadherin knockdown, however, altered cellular geometry, promoting a switch from hepatic to columnar polarity without markedly affecting tubular or primordial BC formation (Fig. 2, D–F). Despite a significant decrease in hepatic polarity, overall apical-basal polarity was maintained in N-cadherin-knockdown cells (Fig. S2 A), indicating that N-cadherin mainly functions to maintain hepatic polarity. Depletion of both E- and N-cadherin in Can 10 cells led to a loss of apical-basal polarity in nearly half the cells (Fig. S2, B and C), a phenotype not observed with individual knockdowns (Fig. S2 A). Notably, some cells failed to maintain cell-cell contacts and detached from clusters (Fig. S2 B). Together, these findings indicate that while E- and N-cadherin cooperate in establishing hepatic polarity and BC formation, presumably by promoting AJ assembly, they play distinct roles in BC elongation and in maintaining hepatic polarity, respectively.

**Figure 2.**
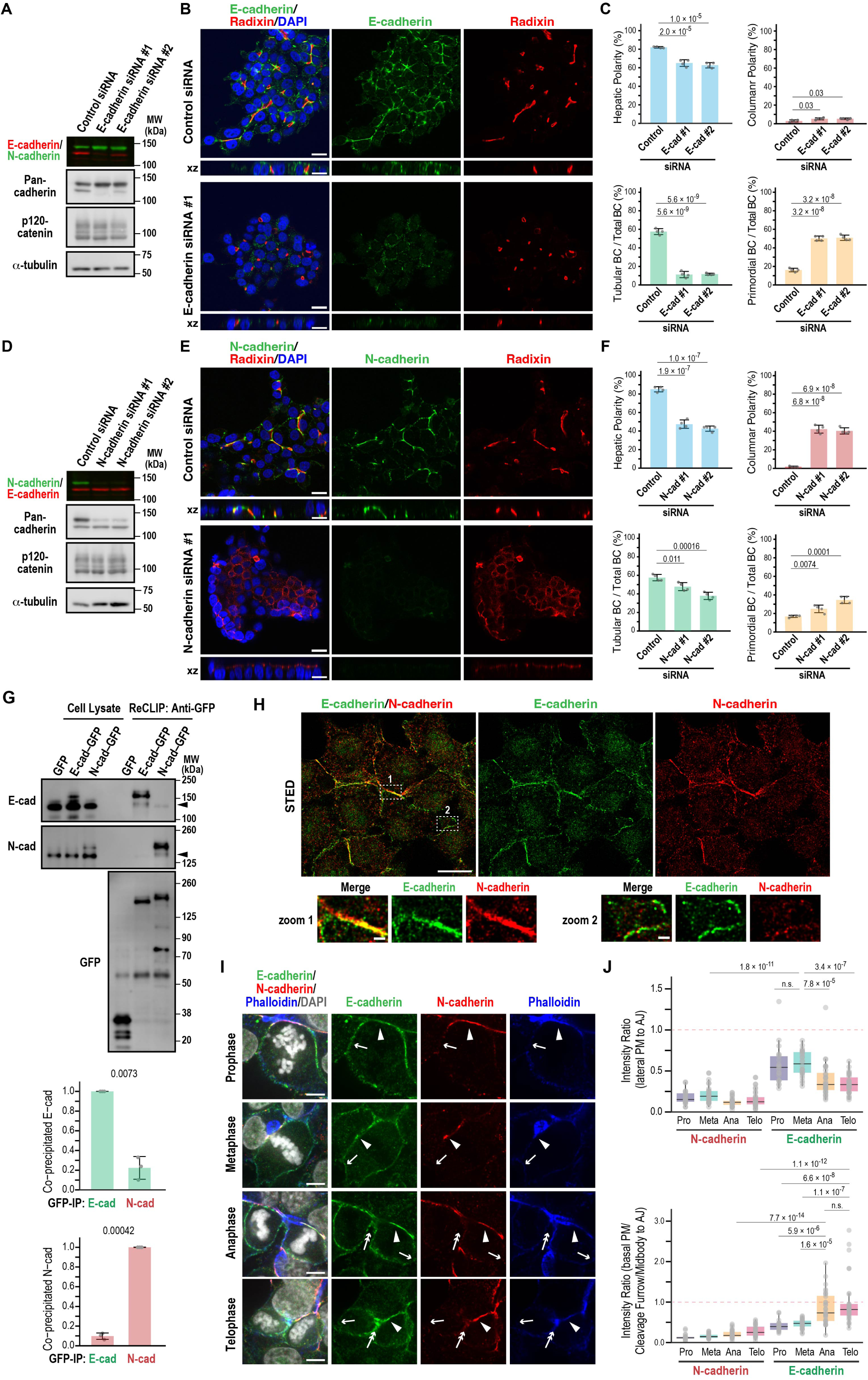
Distinct roles of E- and N-cadherin in BC elongation and hepatic polarity maintenance. (A - C) E-cadherin is required for BC elongation. (A) Immunoblot analysis of Can 10 cell lysates transfected with siRNAs targeting E-cadherin. Molecular weights (MWs) of marker proteins are indicated in kDa. (B) Representative confocal images of Can 10 cells transfected with E-cadherin siRNAs, cultured for 72 h, and stained with DAPI and antibodies against radixin and E-cadherin. The orthogonal views are shown in xz. (C) Quantification of various polarity and BC structures in siRNA-transfected cells. Data represent means ± SD from four independent experiments (≥480 cells per sample). (D - F) N-cadherin is required for maintaining hepatic polarity. Experimental procedures and image analyses were as described in (A – C), except N-cadherin siRNAs were used. Data represent means ± SD from four independent experiments (≥535 cells per sample). (G) E- and N-cadherin preferentially form homodimers or oligomers in Can 10 cells. Top: Lysates and anti-GFP immunoprecipitates from Can 10 cells expressing GFP-tagged E-cadherin or N-cadherin, treated with the reversible crosslinker DSP, were analyzed by immunoblotting using antibodies against E-cadherin, N-cadherin, or GFP. Molecular weights (MWs) of marker proteins are indicated in kDa. Middle and Bottom: Quantification of co-precipitated endogenous cadherins normalized to pulled-down GFP-tagged E-and N-cadherin, respectively. Data represent means ± SD from three independent experiments. (H) Super-resolution imaging of E- and N-cadherin localization. STED microscopy of Can 10 cells cultured for 3 days, fixed, and stained with antibodies against E- and N-cadherin. Lower panels show magnified views of boxed regions. (I) Dynamic localization of E- and N-cadherin during mitosis and cytokinesis. Can 10 cells cultured for 72 h were fixed and stained with phalloidin, DAPI, and antibodies against E- and N-cadherin. Representative confocal images are shown. Arrowheads, arrows, and double-arrows indicate adherens junctions, polar cortex, and the basal cleavage furrow or midbody, respectively. See also Figure S2 D. (J) Quantitative analysis of cadherin distribution at the PM during mitosis and cytokinesis. E- and N-cadherin intensities at lateral (top) or basal (bottom) PM regions were normalized to intensity at AJs. Data are from two-independent experiments (≥ 21 cells per condition). Scale bars, 1 µm (lower panels in H), 5 µm (I), 10 µm (upper panels in H), 20 µm (B and E). p-values are indicated in each graph; n.s., not significant.

The distinct phenotypes resulting from E- versus N-cadherin knockdown raised the possibility that these cadherins predominantly form homo-oligomers. To test this, we transduced Can 10 cells with GFP-tagged E- or N-cadherin via lentiviral infection, and performed reversible cross- link immunoprecipitation (ReCLIP) using the thiol-cleavable cross-linker dithiobis[succinimidyl propionate] (DSP) (Smith et al., 2011). Both GFP-tagged cadherins were expressed at lower levels than their endogenous counterparts (Fig. 2 G). Notably, GFP-tagged cadherins co-precipitated primarily with their respective endogenous forms, though a small amount of E-N hetero-oligomers was also detected (Fig. 2 G). These results support the hypothesis that E- and N-cadherin function mainly as homo-oligomers in Can 10 cells. We next examined cadherin localization using super-resolution stimulated emission depletion (STED) microscopy. E- and N-cadherin co-localized at linear adherens junctions (AJs) (Fig. 2 H, zoom 1), whereas E-cadherin also appeared as puncta or vesicle-like structures near AJs (Fig. 2 H, zoom 1) and at the cell leading edge of cells (Fig. 2 H, zoom 2). A striking difference emerged during cell division (Fig. 2, I and J): N-cadherin remained at AJs surrounding F-actin-decorated BCs (Fig. 2 I, arrowheads), whereas E-cadherin localized both at AJs (arrowheads) and at lateral/polar cortex during pro- and meta-phase (Fig. 2 I, single arrows; and Fig. 2 J). Moreover, E-cadherin was observed at the ingressing basal membrane encompassing the cleavage furrow or midbody during cytokinesis, where new cell-cell contacts form (Fig. 2 I, double arrows; and Fig. 2 J). These observations were validated using multiple independent antibodies (Fig. S2 D). Thus, although cadherins localize to AJs, E-cadherin also appears at the lateral membrane and cleavage furrow during cytokinesis. Taken together, these findings suggest that both E- and N-cadherin function primarily as homo-oligomers contributing to AJ assembly, but diverge in their roles in cell division-linked BC biogenesis (Wang et al., 2014).

### The interplay between E-cadherin and NuMA during cytokinesis is critical for BC elongation

To uncover the molecular mechanisms underlying the distinctive roles of E- and N-cadherin, we performed proximity-dependent biotin identification (BioID) using TurboID-tagged E- and N-cadherin constructs in Can 10 cells (Fig. 3 A; and Fig. S3, A–C) (Cho et al., 2020). Mass spectrometry analysis showed that 74 % (620 out of 834) of the interactome proteins overlapped between E-cadherin and N-cadherin (Fig. S3 C), whereas 16 % (133 out of 834) and 9.7 % (81 out of 834) were specific to E-cadherin and N-cadherin, respectively (Fig. S3 C). Gene ontology (GO) analysis revealed that actin regulators and junction assembly proteins predominated in the overlapping group (Fig. 3 B). This group included the well-known cadherin-binding partners p120-catenin and β-catenin (Fig. 3 A), as well as another interactor, the lipoma preferred partner (LPP) (Van Itallie et al., 2014), confirming the reliability of our BioID assay. Among the E-cadherin-specific proximal proteins, those involved in vesicle transport and those localized at the leading edge were prominent (Fig. 3 B), which is consistent with E-cadherin’s presence at the leading edge (Fig. 2 H; and Fig. S2 B) and at plasma membranes forming cell-cell junctions between daughter cells (Fig. 2 I). Intriguingly, cytokinesis-related proteins were also enriched among the E-cadherin-specific proteins (Fig. 3 B), suggesting a role of E-cadherin in linking cytokinesis to BC elongation. Consistent with our previous report (Wang et al., 2014), confocal time-lapse imaging of Can 10 cells expressing the centrosome marker Mzt1-GFP and the apical marker Radixin-mScarlet revealed that BC elongation is primarily driven by oriented cell division, which enables symmetric inheritance of the pre-existing BC into the daughter cells (Fig. S4, A–C). Among mother cells with a pre- exiting BC, 66% showed a predominantly parallel spindle orientation relative to the BC, leading to symmetric BC inheritance by the daughter cells. In contrast, the remaining 34% exhibited mostly “oblique” or “perpendicular” spindle orientation, resulting in asymmetric BC inheritance. Because of this relationship between oriented cell division and BC elongation, we first focused on Nuclear Mitotic Apparatus protein (NuMA), a nuclear matrix protein that regulates spindle orientation (Kiyomitsu and Boerner, 2021) and forms a complex with E-cadherin (Gloerich et al., 2017) among our identified proximal proteins. NuMA translocates from the nucleus to the cell cortex close to the spindle poles during metaphase and anaphases (Kiyomitsu and Boerner, 2021), suggesting a spatial and temporal overlap with E-cadherin. Strikingly, NuMA knockdown reduced the number of cells with tubular BCs and increased those with primordial BCs (Fig. 3, C–E), with no change in columnar polarity (Fig. 3 E). Cell growth was unaffected at least 72 hours post-siRNA transfection (Fig. S3 D), indicating that the observed defect is not due to a general cytokinesis failure. Given that NuMA depletion phenocopied E-cadherin knockdown, we next examined whether E-cadherin influences NuMA recruitment to the cell cortex during cytokinesis. E-cadherin knockdown did not alter NuMA recruitment to the plasma membrane overall (Fig. 3, F and G; Polar cortex/Cytosol), but specifically impaired its accumulation at the polar cortex, especially during anaphase (Fig. 3, F and G; Polar cortex/Equatorial cortex). These results suggest that E-cadherin contributes to NuMA enrichment at the polar cortex. NuMA cortical localization is mediated by its binding partners—LGN and protein 4.1 family members (4.1R and 4.1G)—which are known to interact with E-cadherin (Gloerich et al., 2017; Yang et al., 2009). Knockdown of these NuMA partners reproduced the BC phenotypes seen with NuMA or E-cadherin depletion (Fig. S3, E–G), supporting the idea that E-cadherin regulates spindle orientation through the NuMA-LGN/4.1 pathway. Consistently, E-cadherin knockdown disrupted mitotic spindle alignment, which in control cells typically runs parallel to the long axis of the pre-existing BC (Fig. S3 H) (Wang et al., 2014). Together, these data indicate that E-cadherin controls BC elongation by regulating spindle orientation via NuMA and its cortical partners. We also explored whether NuMA affects E-cadherin localization. While NuMA knockdown did not impair E-cadherin recruitment to the polar cortex during anaphase (Fig. 3, H and I), it significantly reduced E-cadherin accumulation at the basal region of the cleavage furrow, where new cell-cell contacts form (Fig. 3, H and I). Taken together, these findings suggest that E-cadherin and NuMA functionally cooperate at the polar cortex and cleavage furrow during anaphase and cytokinesis to promote BC elongation.

**Figure 3.**
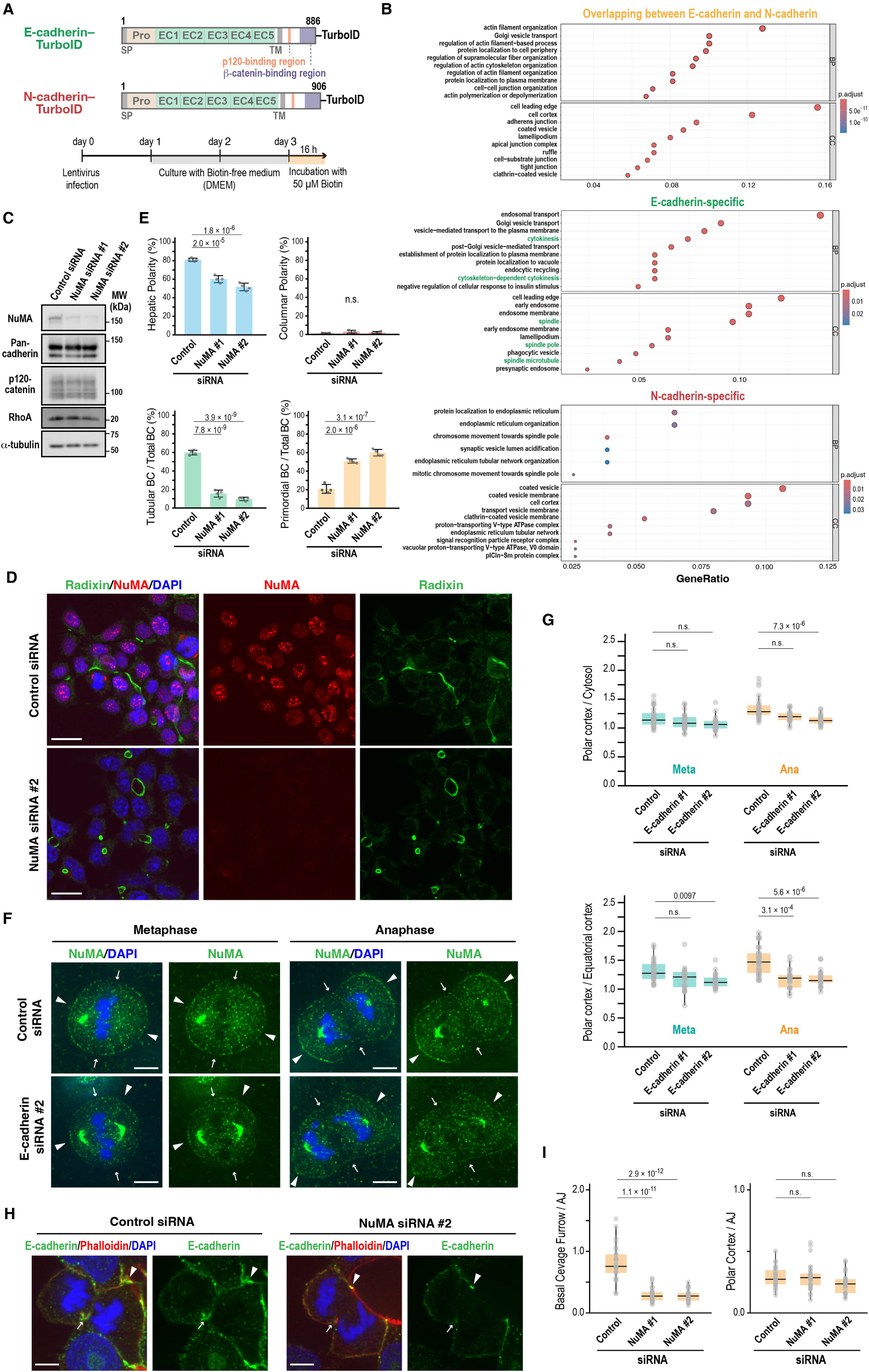
Interplay between E-cadherin and NuMA is critical for BC elongation. (A) Schematic of the BioID assay used to identify proteins in proximity to E- and N-cadherin. (B) Gene Ontology (GO) analysis of E- and N-cadherin-associated proteins. The top 10 terms in BP (Biological Processes) and CC (Cellular Components) are shown. The terms related to spindle and cytokinesis are shown in green. See also Figures S3A-C. (C) NuMA knockdown does not alter expression levels of E-/N-cadherin, p120-catenin, or RhoA. Immunoblot analysis of Can 10 cells transfected with NuMA-targeting siRNAs. Molecular weights (MWs) of marker proteins are indicated in kDa. (D) NuMA is required for BC elongation. Representative confocal images of Can 10 cells transfected with NuMA siRNAs, cultured for 72 h, and stained with DAPI and antibodies against NuMA and radixin. (E) Quantification of polarity and BC features in NuMA-knockdown cells. Data represent means ± SD from four independent experiments (≥836 cells per condition). (F) E-cadherin knockdown disrupts NuMA localization at the polar cortex. Representative confocal images of Can 10 cells transfected with E-cadherin siRNAs, cultured for 72 h, and stained with DAPI and the anti-NuMA antibody. Arrowheads and arrows indicate polar and equatorial cortex, respectively. (G) Quantitative analysis of NuMA localization at the PM during metaphase and anaphase in E-cadherin-knockdown cells. Shown are the ratios of NuMA intensity at the polar cortex to the cytosol (top) and to that at the equatorial cortex (bottom). Data are from two independent experiments (≥21 cells per condition). (H) NuMA knockdown impairs E-cadherin accumulation at the cleavage furrow. Representative confocal images of Can 10 cells transfected with NuMA siRNAs, cultured for 72 h, and stained with DAPI, phalloidin, and the anti-E-cadherin antibody. Arrowheads and arrows indicate adherens junctions and the basal cleavage furrow, respectively. (I) Quantitative analysis of E-cadherin distribution at the PM during anaphase in NuMA-knockdown cells. Ratios of E-cadherin intensity at the basal cleavage furrow (left) or polar cortex (right) to its intensity at AJs are shown. Data are from two independent experiments (≥26 cells per condition). Scale bars, 5 µm (F, H); 20 µm (DE). p-values are indicated in each graph; n.s., not significant.

### RhoA–ROCK functions in BC elongation by recruiting E-cadherin to nascent cell-cell contact sites

The unexpected role of NuMA in E-cadherin localization at the cleavage furrow during anaphase prompted us to further explore its link to cytokinesis. A recent study reported that NuMA restricts the localization of the small GTPase RhoA to the cleavage furrow for efficient cell division in HeLa cells (Sana et al., 2022). We found that in interphase, endogenous RhoA accumulated strongly around the BC area demarcated by ZO-1 in Can 10 cells (Fig. 4 A), but was barely detectable at the plasma membrane in unpolarized Fao cells, despite similar total RhoA levels (Fig. 4 A). To determine the precise spatiotemporal dynamics of RhoA activity, we expressed a probe specific for its active (GTP-bound) form—dimeric Tomato-tagged tandem repeats of rhotekin GBD (dT-2xrGBD)—in Can 10 cells (Mahlandt et al., 2021). Importantly, this probe did not affect hepatic polarization or BC development (Fig. 4 B). Consistent with the antibody staining (Fig. 4 A), the dT-2xrGBD probe showed strong signal intensity surrounding the BC—appearing as a dark space between adjacent cells (Fig. 4 C, time point -50). As the cell progressed through cytokinesis, the probe signal intensified at the BC membrane of the dividing cell, peaking as a punctum at the division site (Fig. 4 C, time point -10; and Fig. 4 D). This punctum likely represents the midbody between daughter cells (Hu et al., 2012). The intensity gradually declined following cleavage furrow ingression (Fig. 4 C, time point 0). This dynamic pattern suggests that active RhoA may contribute to BC maintenance and elongation during cytokinesis. To investigate this hypothesis, we depleted RhoA using siRNAs (Fig. 4 E). Knockdown cells failed to form tubular BCs and instead displayed an increased number of primordial BCs, without an increase in columnar polarity (Fig. 4, F and G). Since ROCK2 was the only E-cadherin-specific proximal protein identified among known RhoA downstream effectors (Fig. S5 A), we examined its role in BC morphogenesis. Similar to RhoA depletion, treatment of Can 10 cells with the ROCK inhibitor Y27632 significantly reduced tubular BC formation (Fig. 4, H and I). This phenotype was corroborated by *ex vivo* treatment of mouse fetal liver with Y27632, which impaired BC development (Fig. 4 J). Together, these results indicate that the RhoA–ROCK pathway is essential for proper BC development. Given the phenotypic similarities between RhoA and NuMA knockdowns in Can 10 cells and their functional connection during cytokinesis in other cell types (Sana et al., 2022), we examined the effect of NuMA depletion on RhoA localization during anaphase. As expected, NuMA knockdown reduced RhoA recruitment to the basal side of the cleavage furrow (Fig. 4, K and L). Considering the role of NuMA in E-cadherin localization at the cleavage furrow (Fig. 3, H and I), these findings raise the possibility that NuMA may regulate this E-cadherin localization by restricting RhoA–ROCK to the cleavage furrow. To test this, we treated cells with Y27632 and then examined E-cadherin localization during anaphase and telophase. Notably, this treatment reduced E-cadherin recruitment to the daughter cell interface at both stages, without affecting N-cadherin localization (Fig. 4, M and N). These results suggest that the ROCK kinase activity is required for E-cadherin recruitment at the newly forming cell-cell contacts, facilitating the sealing and elongation of a pre-existing BC.

**Figure 4.**
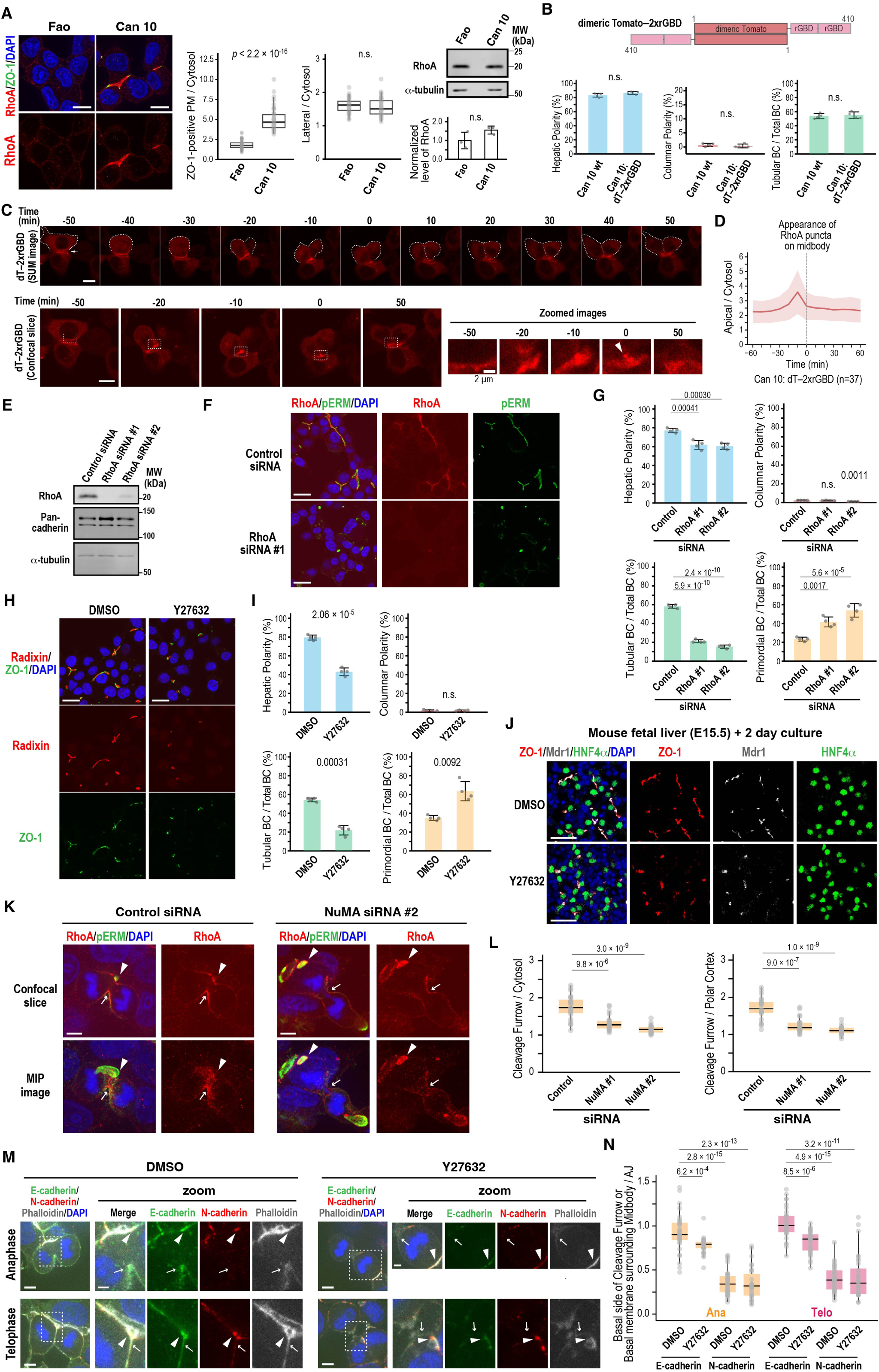
RhoA–ROCK signaling regulates E-cadherin localization during cytokinesis and drives BC elongation. (A) RhoA accumulates at the BC region in polarized Can 10 cells but not in unpolarized parental Fao cells. Left: Representative confocal images of Can 10 and Fao cells cultured for 3 days and stained with DAPI and antibodies against RhoA and ZO-1. Middle: Quantification of RhoA intensity at ZO-1-positive plasma membranes (left) or lateral membranes (right), normalized to cytoplasmic intensity. Values are from two independent experiments (≥50 cells per condition). Right: Immunoblot analysis and quantification of RhoA levels in Fao and Can 10 cells. Data represent means ± SD from three independent experiments; RhoA/α-tubulin ratio in Fao cells is normalized to 1.0. (B) Expression of an active RhoA biosensor (dTomato–2×rGBD) does not impair polarization or BC formation in Can 10 cells. Top: schematic of the dTomato–2×rGBD construct. Bottom: Quantification of Can 10 cells expressing the biosensor, categorized by polarity type and BC morphology. Data represent means ± SD from four independent experiments (≥268 cells per condition). (C - D) The RhoA biosensor accumulates at the apical region and cleavage furrow during cytokinesis in cells with a pre-existing BC. (C) Montage of time-lapse imaging from Can 10 cells expressing dT–2×rGBD during cytokinesis. SUM projection and confocal images are shown. Zoomed views of the boxed regions in the confocal images are also shown (bottom right). The arrow in the SUM image at time -50 indicates a pre-existing BC; the arrowhead in the zoomed image at time 0 indicates the midbody. (D) Quantification of dT–2×rGBD intensity at the BC region over time from cells in (C) (n =37) were quantified and plotted. Bold lines and shaded bands represent the mean and SD, respectively, as used throughout this study. (E - G) RhoA is required for BC elongation. (E) Immunoblot analysis of RhoA-knockdown Can 10 cells. Molecular weights (MWs) of marker proteins are indicated in kDa. (F) Representative confocal images of RhoA-depleted Can 10 cells stained with DAPI and antibodies against RhoA and phospho-ezrin/radixin/moesin (pERM). (G) Quantification of polarity and BC features in RhoA-knockdown cells. Data represent means ± SD from four independent experiments (≥593 cells per condition). (H - I) ROCK activity is required for BC elongation. (H) Confocal images of Can 10 cells treated with the ROCK inhibitor Y27632. Treatment was initiated one day after plating at a concentration of 3 µM and was continued for 48 h. Cells were stained with DAPI and antibodies against ZO-1 and radixin. (I) Quantification of polarity and BC features in Y27632-treated cells. Data represent means ± SD from four independent experiments (≥666 cells per condition). (J) ROCK activity is required for BC elongation in ex vivo mouse livers. Liver lobes from E15.5 were treated with 1 µM Y27632 for two days and stained with antibodies against ZO-1, Mdr1, and HNF4α for confocal analysis. (K - L) NuMA is required for RhoA accumulation at the cleavage furrow. (K) Confocal images (top) and maximum intensity projection (MIP) images of Can 10 cells transfected with NuMA siRNAs, stained with DAPI and antibodies against RhoA and phospho-ERM. Arrowheads and arrows indicate BCs and the basal cleavage furrow, respectively. (L) Quantification of RhoA intensity at the basal cleavage furrow relative to cytoplasm (left) and polar cortex (right). Data are from two-independent experiments (≥22 cells per condition). (M - N) ROCK activity is required for E-cadherin accumulation at the division site. (M) Representative confocal images of DMSO- or Y27632-treated cells stained with DAPI, phalloidin, and antibodies against E-cadherin and N-cadherin. Arrowheads and arrows indicate adherens junctions and the basal cleavage furrow or midbody, respectively. (N) Quantification of E-cadherin intensity at the basal cleavage furrow and midbody relative to AJs. Data are from three-independent experiments (≥30 cells per condition). Scale bars, 2 µm (zoomed images in C and M), 5 µm (K, M), 10 µm (A, C), 20 µm (F, H); 50 µm (J). p-values are indicated in each graph; n.s., not significant.

### ARHGEF17 is the Rho GEF responsible for BC elongation

Since RhoA is activated at the apical region (presumably at the AJ) and the cleavage furrow of dividing cells during BC elongation, we aimed to identify the guanine nucleotide exchange factors (GEFs) responsible for these activations. Among the RhoA-specific GEFs captured by our BioID assay, ARHGEF17/TEM4 is the only one reported to function in both AJ assembly (Ngok et al., 2013) and cytokinesis (Prifti et al., 2025), although its localization at the cleavage furrow has not been described but is expected (Fig. 5 A; and Fig. S5 B). Thus, we first examined its localization in Can 10 cells. ARHGEF17–GFP colocalized with E-cadherin around BCs (Fig. 5 B). This localization was reduced following treatment with Latrunculin A (LatA), an F-actin depolymerization agent (Fig. S5, C and D) (Spector et al., 1989; Yarmola et al., 2000). A combined treatment with LatA and E-cadherin knockdown abolished this localization (Fig. S5, D and E), suggesting that ARHGEF17 requires both F-actin and E-cadherin for its recruitment to AJs. This is consistent with a previous report that ARHGEF17 localizes to cell-cell contacts in a cadherin-catenin complex-dependent manner (Ngok et al., 2013). To examine the dynamics of ARHGEF17 during cytokinesis, we tracked ARHGEF17– GFP with the mScarlet-tagged cytokinesis marker Myl12b, an isoform of myosin regulatory light chains (Brito and Sousa, 2020). Live cell imaging revealed that ARHGEF17 localized to the cleavage furrow and then concentrated around the midbody (Fig. 5 C, time points 40 and 80). Interestingly, when the long axis of a dividing cell was aligned parallel to a pre-existing BC, its midbody formed near the BC (Fig. 5 C, Cell #1). In contrast, when the long axis of a dividing cell was oblique to the BC, its midbody was initially formed away from the BC but was eventually pulled and incorporated into the pre-existing BC (Fig. 5 C, Cell #2). Higher magnification image analysis showed that ARHGEF17 was distributed at the daughter cell interface around the midbody, where E-cadherin was also observed (Fig. 5 D). These findings suggest that ARHGEF17 plays a role in coordinating cytokinesis with new cell-cell contact formation.

**Figure 5.**
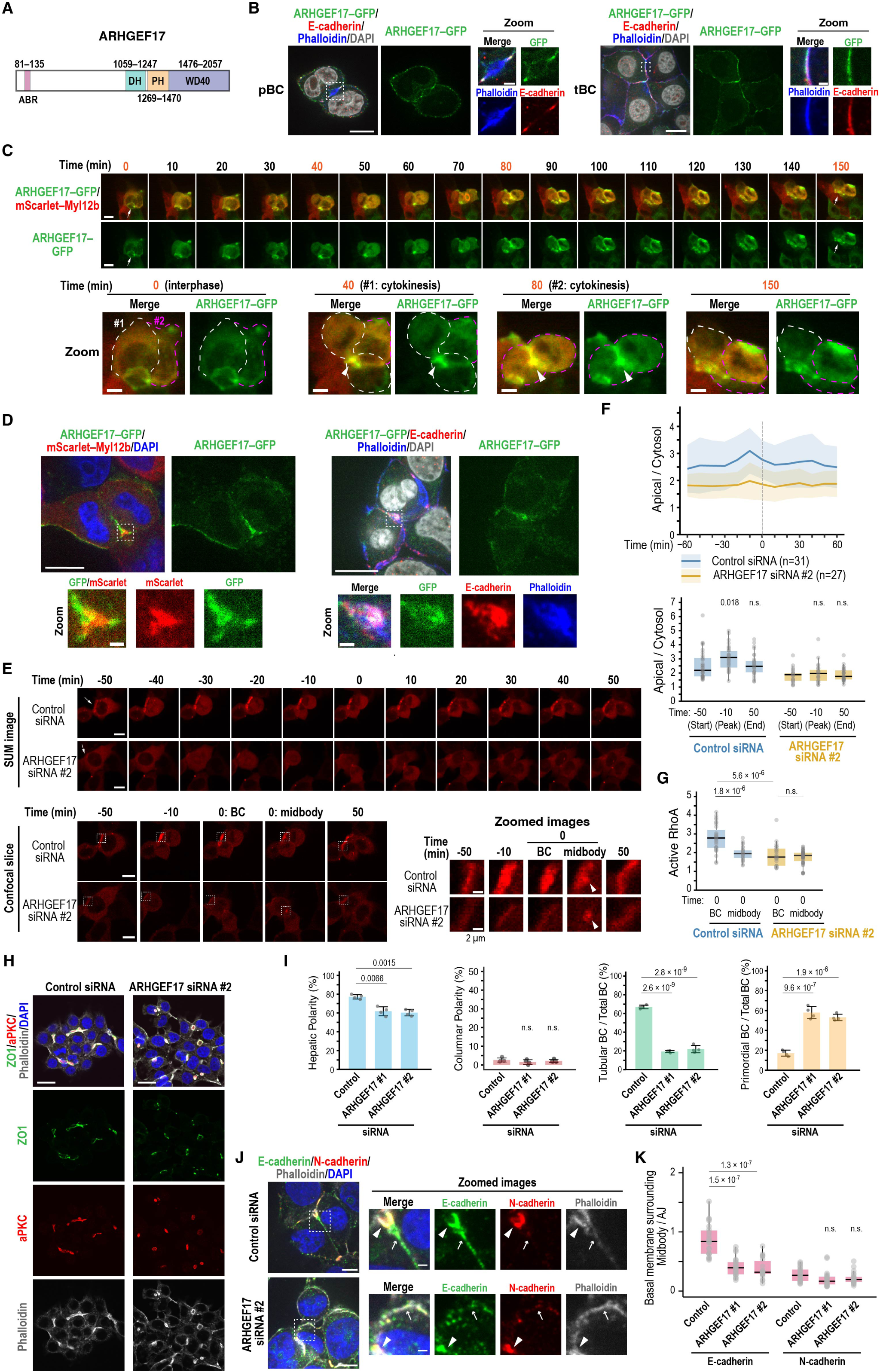
ARHGEF17 activates RhoA at the apical region and cleavage furrow to promote BC elongation. (A) Schematic of rat ARHGEF17. ABR, actin-binding region; DH, Dbl homology domain; PH, Pleckstrin homology domain, WD40, WD40-repeat domain. (B) ARHGEF17 localizes to the cell cortex and AJs in cells with primordial or tubular BCs. Representative confocal images of Can 10 cells expressing ARHGEF17–GFP, cultured for 2 days post-transduction, and stained with DAPI, phalloidin, and anti-E-cadherin antibody. (C) ARHGEF17 accumulates at the apical region and cleavage furrow during cytokinesis. Montage of time-lapse imaging of Can 10 cells expressing ARHGEF17–GFP and mScarlet– Myl12b during cytokinesis. SUM projections at time 0, 40, 80, and 150 are shown at the bottom. Cells #1 and #2 are outlined in white and magenta, respectively. (D) ARHGEF17 localizes to the daughter cell interface near the midbody during late cytokinesis. Representative confocal images of Can 10 cells expressing ARHGEF17–GFP alone (right) or with mScarlet–Myl12b (left) cultured for 2 days post-transduction, and stained as indicated. (E) ARHGEF17 is required for RhoA activation at the apical region and cleavage furrow during cytokinesis. Time-lapse montage of Can 10 cells transfected with control or ARHGEF17 siRNAs and expressing dT–2×rGBD. SUM projects (top), confocal images (bottom), and zoomed views of boxed regions are shown. Arrows (-50 min) indicate BCs; arrowheads (0 min) indicate the midbody. (F) Quantitative analysis of RhoA activation in cells with a pre-existing BC. Ratios of dT– 2×rGBD intensities at the BC vs. cytoplasm in control and ARFGEF17-knockdown cells imaged in (E) are plotted over time (top), and at selected timepoints (bottom). (G) ARHGEF17 primarily activates RhoA at the apical region, but not at the midbody. Ratios of dT–2×rGBD intensity at the BC or midbody to cytoplasm are shown in cells from (E). (H) ARHGEF17 is required for BC elongation. Representative confocal images of Can 10 cells transfected with control siRNA or ARHGEF17 siRNA, cultured for 72 h, and stained with DAPI, phalloidin, and antibodies against aPKC and ZO-1. (I) Quantification of polarity and BC structures in control and ARHGEF17-knockdown Can 10 cells. Data represent means ± SD from four independent experiments (≥373 cells per condition). (J) ARHGEF17 is required for E-cadherin accumulation at the cleavage furrow adjacent to the midbody. Confocal images of control and ARHGEF17-knockdown cells stained with DAPI, phalloidin, and antibodies against E- and N-cadherin. Arrowheads and arrows indicate adherens junctions and the basal midbody, respectively. (K) Quantitative analysis of E- and N-distribution at the division site. Ratios of cadherin intensity at the basal side of the midbody to their intensity at AJs are shown in cells from (J). Data are from two-independent experiments (≥21 cells per condition). Scale bars, 1 µm (zoomed images in B, D, and J), 2 µm (zoomed images in E), 5 µm (lower images in C), 10 µm (B, D, E, and upper images in C), 20 µm (H). p-values are shown above each graph; n.s., not significant.

To assess whether ARHGEF17 activates RhoA during cytokinesis-linked BC elongation, we analyzed dT-2xrGBD-expressing cells with or without ARHGEF17 knockdown (Fig. 5 E; and S5, F–H). In contrast to control cells, which showed high RhoA activity around the BC and its elevation during cytokinesis, ARHGEF17 knockdown cells exhibited decreased basal RhoA activity on the BC (Fig. 5 E; and Fig. 5 F, time point -50) and failed to increase RhoA activity during cytokinesis (Fig. 5, E and F). Furthermore, ARHGEF17 depletion led to increased asymmetric BC inheritance (Fig. S5 H), suggesting a role in the regulation of spindle orientation. However, RhoA intensity at the midbody was not significantly affected in knockdown cells, suggesting that other GEF(s) may be responsible for activating RhoA at this site (Fig. 5 G). These results indicate that ARHGEF17 is essential for RhoA activation around BCs especially during cytokinesis, linking cytokinesis to BC development. In support of this, ARHGEF17 knockdown impaired tubular BC formation (Fig. 5, H and I) but increased primordial BCs, phenocopying RhoA knockdown and ROCK inhibition (Fig. 4, G and I). We also tested whether ARHGEF17 contributes to cadherin localization at later stages of cytokinesis, and found that ARHGEF17 depletion impaired E-cadherin recruitment to the daughter cell interface during telophase (Fig. 5, J and K). Thus, ARHGEF17 is required for RhoA activation and E-cadherin accumulation and promotes cytokinesis-linked BC elongation.

### p190B and ARVCF mediate the role of N-cadherin in maintaining hepatic polarity

We next sought to understand how N-cadherin deficiency induces columnar polarity. Previous studies suggest that elevated RhoA activity alters polarity orientation in 3D-cultrued epithelial cells (Yu et al., 2008) and is associated with columnar but not hepatic polarity (Lazaro-Dieguez and Musch, 2017). This raises the possibility that N-cadherin maintains hepatic polarity by inactivating RhoA activated by E-cadherin and ARHGEF17 in a spatiotemporally controlled manner. Thus, we searched our BioID data for Rho GTPase-activating proteins (RhoGAPs). Notably, only two RhoA-specific GAPs were identified: ARHGAP5/p190B and ARHGAP35/p190A, with p190B specifically found as an N-cadherin proximal protein (Fig. 6 A). Consistent with this, p190B knockdown caused a significant shift from hepatic to columnar polarity, whereas p190A depletion had no effect (Fig. 6, B and C; and S6 A). Double knockdown of both p190A and p190B did not further enhance columnar polarity compared to p190B single depletion alone, suggesting no redundant roles in this process (Fig. 6, B and C; and S6 A). These findings indicate that p190B-mediated RhoA inactivation is essential for maintaining hepatic polarity.

**Figure 6.**
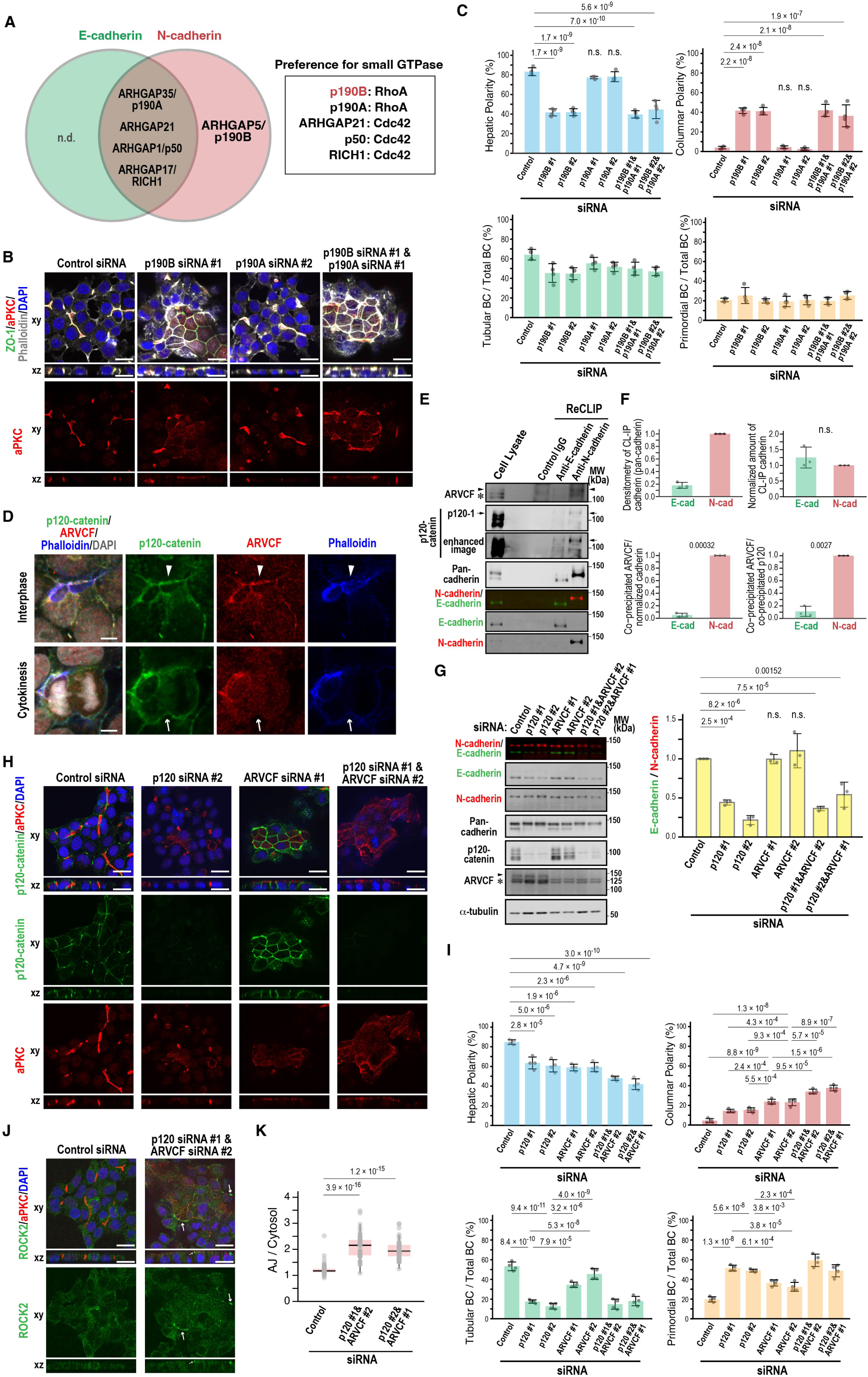
p190B and ARVCF mediate the role of N-cadherin in hepatic polarity maintenance by downregulating RhoA activity. (A) Venn diagram of ARHGAP proteins identified by BioID analysis. Their preferences for Rho family small GTPases are shown on the right. The RhoGAP specific to N-cadherin is indicated in red. Note that E-cadherin-specific group does not include any ARHGAPs. n.d., not detected. (B) p190B, but not p190A, is required for hepatic polarity maintenance. Representative confocal images of Can 10 cells transfected with p190A and/or p190B siRNAs, cultured for 72 h, and stained with DAPI and antibodies against ZO-1 and aPKC. See also Figure S6 A. (C) Quantification of polarity and BC structures in cells from (B). Data represent means ± SD from four independent experiments (≥431 cells per condition). (D) Subcellular localization of p120-catenin and ARVCF during interphase and cytokinesis. Representative iSIM images of Can 10 cells cultured for 3 days and stained with DAPI, phalloidin, and antibodies against p120-catenin and ARVCF. (E) N-cadherin, but not E-cadherin, preferentially associates with ARVCF over p120-catenin. Lysates of Can 10 cells treated with the reversible crosslinker DSP were immunoprecipitated using control IgG, anti-E-cadherin or anti-N-cadherin antibody, followed by SDS-PAGE and immunoblot analysis. Molecular weights (MWs) of marker proteins are indicated in kDa. (F) Quantification of ARCVF and p120-catenin binding preference for E- and N-cadherin. Top: Intensities of pulled-down cadherins (left) and their normalized protein amounts (right). Bottom: Ratios of co-precipitated ARVCF (left) or ARVCF/p120-catenin (right) normalized to the cadherin pulldown. Data represent means ± S.D. from three independent experiments. The ratio of ARVCF to N-cadherin (left) and ARVCF vs. p120-catenin to N-cadherin (right) is set to 1.0. See also Figure S1 C. (G) Knockdown of p120, but not ARVCF, destabilizes E-cadherin without affecting N- cadherin. Left: Representative immunoblot blots of single and double knockdown cells. Right: Quantification of E-cadherin to N-cadherin ratios. Data represent means ± SD from three independent experiments; the ratio in control siRNA-transfected cells is set to 1.0. (H) ARVCF primarily regulates hepatic polarity, while p120-catenin affects BC elongation. Representative confocal images of Can 10 cells transfected with ARVCF and/or p120-catenin siRNAs, cultured for 72 h, and stained with DAPI and antibodies against p120-catenin and aPKC. (I) Quantification of polarity and BC structures in cells from (F). Data represent means ± SD from four independent experiments (≥803 cells per condition). (J) ROCK2 accumulates at AJs in ARVCF/p120-catenin double knockdown cells. Representative confocal images of Can 10 cells transfected with both siRNAs, cultured for 72 h, and stained with DAPI and antibodies against aPKC and ROCK2. Arrows indicate ROCK2 accumulation on the apical most junctions. (K) Quantification of ROCK2 localization at AJs. Ratios of ROCK2 intensity at the apical edge vs. cytoplasm are shown for control and double knockdown cells. Data are from two-independent experiments (≥51 cells per condition). Scale bars, 5 µm (D), 20 µm (B, H, J). p-values are indicated at the top of each graph; n.s., not significant.

Since p190B interacts with p120-catenin and its paralog ARVCF (Cho et al., 2010; Ponik et al., 2013; Wildenberg et al., 2006), we investigated the localization and function of these catenins in Can 10 cells. While p120-catenin and ARVCF co-localized at AJs during interphase (Fig. 6 D), but showed distinct localization during cytokinesis: p120-catenin was present at the plasma membrane, including the cleavage furrow, whereas ARVCF was restricted to AJs around BCs (Fig. 6 D, cytokinesis), reminiscent of the differential localization of E-cadherin and N-cadherin (Fig. 2 I). Consistent with this, ReCLIP assays using DSP demonstrated that endogenous ARVCF was co-precipitated more efficiently with N-cadherin than with E-cadherin (Fig. 6, E and F; and Fig. S1, C and D). ARVCF also showed a preference for N-cadherin over p120-catenin (Fig. 6 E). The association between p120-catenin and E-cadherin was confirmed by the observation that only E-cadherin was markedly reduced in p120-catenin knockdown cells, whereas ARVCF depletion did not affect cadherin stability (Fig. 6 G). These results suggest that p120-catenin and ARVCF preferentially interact with E-cadherin and N-cadherin, respectively, with p120-catenin specifically stabilizing E-cadherin. To assess their roles in hepatic polarity and BC development, we examined single and double knockdowns of p120-catenin and ARVCF (Fig. 6 G). p120-catenin knockdown decreased tubular BCs but increased primordial BCs (Fig. 6, H and I), similar to E-cadherin depletion. In contrast, ARVCF knockdown had little effect on tubular BC formation but increased columnar polarity (Fig. 6, H and I). Notably, columnar polarity was further enhanced by combined depletion of p120-catenin and ARVCF (Fig. 6, H and I). These results suggest that p120-catenin primarily functions with E-cadherin in BC elongation, while ARVCF, with minor input from p120- catenin, functions with N-cadherin to maintain hepatic polarity.

We next asked whether RhoA signaling was activated in columnar cells induced by N-cadherin depletion or by double knockdown of p120-catenin and ARVCF. In control cells, ROCK2 was predominantly cytoplasmic and barely detectable at the plasma membrane, whereas in N-cadherin-depleted cells with columnar polarity it concentrated at the apical side of cell-cell contacts (Fig. S6, B and C). A similar pattern was observed in columnar cells induced by the combined knockdown of p120-catenin and ARVCF (Fig. 6, J and K). These findings suggest that aberrant activation of RhoA–ROCK occurs upon N-cadherin loss or double depletion of p120-catenin and ARVCF, likely due to the absence and/or inactivation of p190B at AJs. These findings suggest that ARVCF and its binding partner p190B mediate the role of N-cadherin in maintaining hepatic polarity by inactivating RhoA at AJs.

## Discussion

In this study, we reveal that hepatocytes employ E- and N-cadherin to regulate hepatic polarity and drive BC biogenesis through spatiotemporal control of RhoA activity in an opposing manner, addressing a fundamental question in liver biology. While both cadherins cooperate in polarity establishment, they also serve distinct, non-redundant roles. E-cadherin drives cytokinesis-linked BC elongation by coordinating spindle orientation and RhoA activation through NuMA and the RhoGEF ARHGEF17, which also facilitates its recruitment to daughter cell interfaces for nascent cell-cell contact formation. In contrast, N-cadherin maintains hepatic polarity by promoting RhoA inactivation via ARVCF and the RhoGAP p190B. Collectively, these findings define the shared and unique roles of E- and N-cadherin in hepatic polarity and BC formation, processes essential for liver architecture and function.

### Co-expression of E- and N-cadherin and ECM signaling are required to establish hepatic polarity

What dictates hepatic polarity? A clue emerged from our single-cell transcriptomic analyses, which revealed that bipotential hepatoblasts follow “default” and “directed/branched” pathways to differentiate into hepatocytes and cholangiocytes, respectively (Wang et al., 2020; Yang et al., 2017; Yang et al., 2023). This divergence occurs between E13.5 and E15.5 (Wang et al., 2020; Yang et al., 2017; Yang et al., 2023). In both mouse and human, hepatocyte differentiation is accompanied by specific changes in histone modifications, suggesting a potential shared intermediate stage between hepatocytes and neurons prior to terminal differentiation (Yang et al., 2023).

Neurons predominantly express N-cadherin, which mediates both adherens junctions between neural progenitor cells during development (Miyamoto et al., 2015; Punovuori et al., 2021; Radice, 2013; Taneyhill, 2008) and trans-cellular adhesion at the synaptic clefts for neuronal transmission (Tanaka et al., 2000). In line with this, hepatoblasts and hepatocytes express both E-cadherin, reflecting their epithelial nature, and N-cadherin, reflecting neuronal similarity, during BC formation and elongation between E13.5 and E17.5 (Doi et al., 2007). Strikingly, when the liver lobules are established postnatally, cadherin distribution becomes zonally restricted: E-cadherin is predominantly expressed in the periportal vein region (Zone 1), whereas N-cadherin shows a complementary pattern with highest expression in the peri-central vein region (Zone 3) (Doi et al., 2007). During regeneration, hepatocytes, especially those from the intermediate region (Zone 2), can upregulate E-cadherin, resulting in co-expression of both cadherins (He et al., 2021; Lin et al., 2023; Wang et al., 2024). These findings suggest that E- and N-cadherin co-expression contributes to the establishment of hepatic polarity and BC biogenesis, a possibility supported by our observations. They also indicate that Can 10 cells resemble developing and regenerating hepatocytes, rather than adult hepatocytes.

In addition to E- and N-cadherin co-expression, extracellular matrix (ECM) signaling plays a decisive role in establishing hepatic polarity. Hepatocytes and cholangiocytes are associated with distinct ECM environments. The basal side of cholangiocytes rests on a dense basal basement rich in network-forming collagen IV and laminin (Tanimizu and Mitaka, 2017), whereas the basal side of hepatocytes faces the sinusoidal space, where only a sparse extracellular milieu composed of collagen fibers, fibronectin, and proteoglycans is present (Tanimizu and Mitaka, 2017; Treyer and Musch, 2013). This stark difference in ECM composition likely contributes to hepatic versus columnar polarity (Treyer and Musch, 2013). Supporting this notion, the biopotential liver progenitor cell line HPPL adopts cholangiocyte-like polarity when cultured in Matrigel (a basement membrane extract) (Tanimizu et al., 2007; Tanimizu et al., 2004). Similarly, hepatoblasts from E11.5—E14.5 embryos adopt hepatocyte or cholangiocyte fates when induced by gelatin (a collagen extract) or Matrigel, respectively (Lee et al., 2016; Lv et al., 2015; Wang et al., 2020). Remarkably, we observed that Can 10 cells cultured on Matrigel lost hepatic polarity and formed multi-cellular clusters surrounding a central lumen, reminiscent of bile duct-like structures, which is in sharp contrast to the Can 10 cells cultured on collagen I, displaying hepatic polarity and forming BC structures (Fig. S6 D). This highlights a striking similarity between Can 10 cells and bipotential hepatoblasts. The mechanisms underlying this ECM-instructed choice between hepatic polarity/BC structures and columnar polarity/duct-like structures remain to be elucidated.

Collectively, our findings indicate that E- and N-cadherin co-expression together with specialized ECM signaling are required for the establishment of hepatic polarity, which is critical for liver tissue architecture and function.

### Opposing regulation of RhoA activity by E- and N-cadherin enables BC elongation and hepatic polarity

E- and N-cadherin are known to alter their expression and play distinct roles in processes such as EMT (Loh et al., 2019; van Roy, 2014), but the underlying mechanisms have remained elusive. Taking advantage of their unique co-expression in hepatocytes and using the BioID assay, we identified specific interactors and found that E- and N-cadherin regulate hepatic polarity and BC biogenesis by exerting opposite control on RhoA activity (Fig. 7 A).

**Figure 7.**
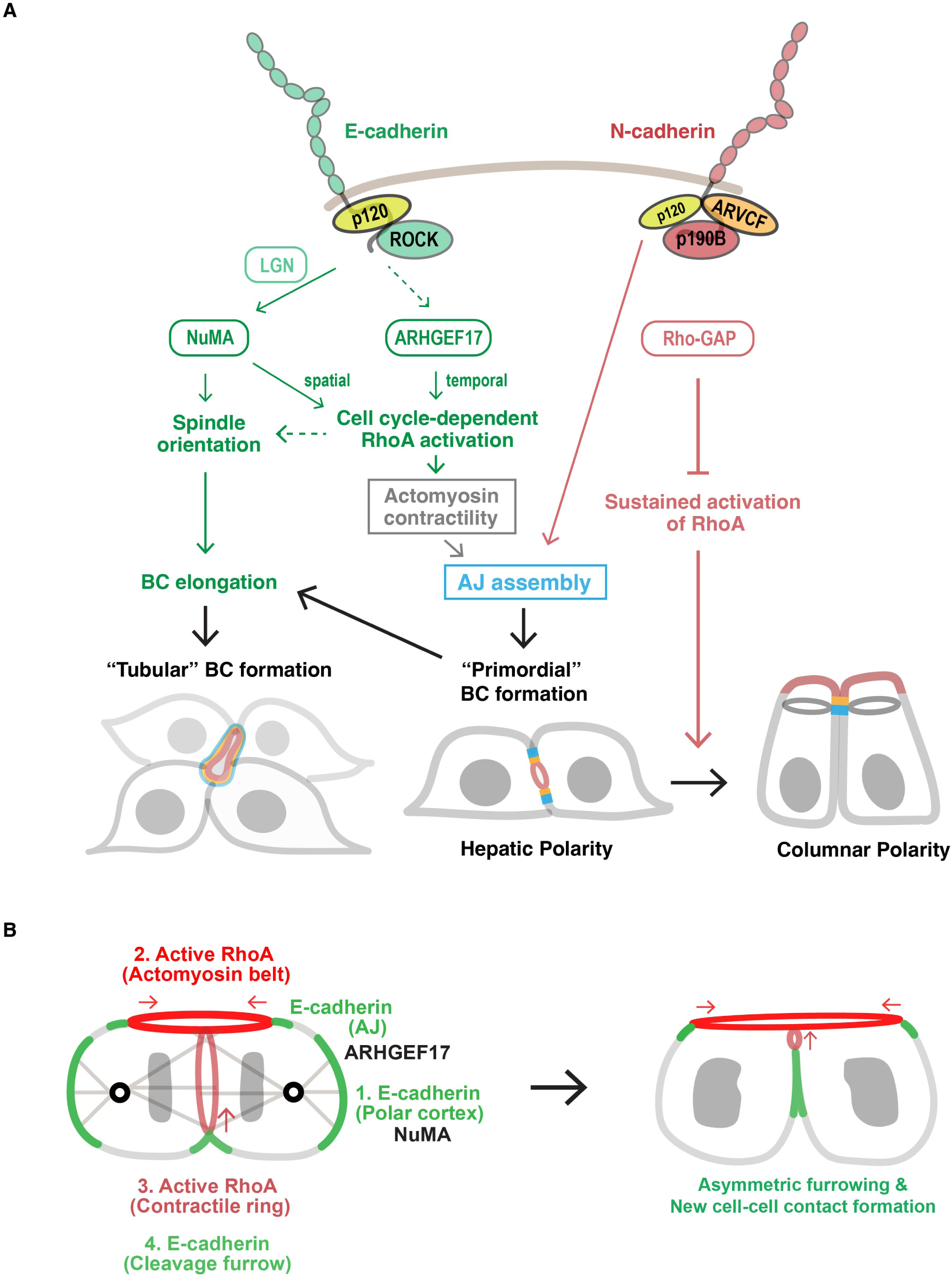
Models for cytokinesis-linked hepatocyte polarization and BC formation. (A) Schematic illustrating the overlapping and distinct roles of E-cadherin and N-cadherin in controlling BC elongation and maintaining hepatic polarity via opposing effects on RhoA activity. E-cadherin activates RhoA at the AJs surrounding the BC and at the cleavage furrow via ARHGEF17, while restricting active RhoA to the division site via NuMA at the lateral membrane during cytokinesis. This spatially restricted RhoA activity enables the recruitment of E-cadherin to nascent cell-cell contacts between daughter cells, promoting new AJ assembly during primordial BC formation and BC elongation. ARHGEF17 and NuMA also contribute to BC elongation by regulating spindle orientation (see below). While N-cadherin shares the role of promoting AJ assembly during primordial BC formation with E-cadherin, it also decreases RhoA activity at AJs via ARVCF and p190B RhoGAP, thereby preventing an unwanted switch from hepatic to columnar polarity. (B) Model depicting the roles of two distinct mechanical forces in regulating spindle orientation during BC elongation. The first force is generated by E-cadherin—NuMA— dynein—astral microtubule pathway at the cell poles (#1), which governs the initial selection of spindle orientation. The second force arises from a RhoA-mediated actomyosin belt at the AJs surrounding the BC membrane, activated by ARHGEF17 (#2). This force stabilizes spindle orientation by anchoring the dividing cell at the BC side of its cleavage furrow and also provides an asymmetric mechanical cue that biases furrow ingression towards the BC (#3 and red arrow). RhoA–ROCK-dependent constriction of the contractile ring enables E-cadherin delivery to the division site during cytokinesis, promoting the formation of a new AJ between daughter cells and driving BC elongation (#4).

E-cadherin acts through its proximal binding partners, NuMA and ARHGEF17, to regulate spindle orientation and RhoA activation, promoting cytokinesis-linked BC elongation. Time-lapse analysis revealed that among dividing cells with a pre-existing BC, ∼66% display symmetric BC inheritance with elongation, most dividing with a spindle parallel to the BC long axis. The remaining ∼34% display asymmetric inheritance without elongation, most dividing with an oblique or perpendicular spindle relative to the BC (Fig. S4). Based on these observations and the temporal sequence of BC morphogenesis during liver development (Fig. 1 A; and Fig. S1 C), we hypothesize that spindle orientation is developmentally regulated, with more parallel divisions driving BC elongation during early development (E15.5—E17.5) and more oblique/perpendicular divisions promoting hepatocyte stratification into cord-like structures and BC network formation during late development (P0, perinatal stage).

Our findings also suggest a mechanistic model in which two mechanical forces coordinate to dictate spindle orientation in E-cadherin-mediated parallel divisions during BC elongation (Fig. 7 B). The first force acts at the cell poles, mediated by the E-cadherin—NuMA—dynein— astral microtubules (MTs) pathway (di Pietro et al., 2016; Gloerich et al., 2017; Kiyomitsu and Boerner, 2021) and is responsible for the initial selection of spindle orientation (step 1). The second force acts at AJs surrounding the BC, generated by the ARHGEF17- and RhoA-dependent actomyosin belt, and stabilizes spindle orientation (step 2). This contractile activity also provides an asymmetric mechanical cue that biases furrow ingression towards the BC via the RhoA-activated cytokinetic actomyosin ring (step 3), a process common to epithelial planar divisions (Fleming et al., 2007; Jinguji and Ishikawa, 1992; Maddox et al., 2007; Thieleke-Matos et al., 2017; Wang et al., 2014). Following cytokinesis, E-cadherin delivery to the division site enables new AJ assembly between daughter cells (step 4).

This model predicts that disrupting either force—actomyosin at AJs or MT/dynein at the poles—impairs spindle orientation and thereby inhibits BC elongation. Supporting this, depletion of E-cadherin, NuMA, ARHGEF17, or RhoA, or inhibition of ROCK blocked BC elongation (Fig. 2–5). Mechanistically, E-cadherin depletion reduced spindle orientation, partly by decreasing NuMA localization at the polar cortex during anaphase (Fig. 3, F and G; and Fig. S3 H), while ARHGEF17 depletion impaired RhoA activation at AJs, also altering spindle orientation (Fig. 5, E–G; and Fig. S5 H). Notably, an AJ-associated spindle orientation mechanism (“rescue mechanism”) had been proposed in columnar epithelial cells such as MDCK cells but thought absent in hepatocytes (Lazaro-Dieguez and Musch, 2017). Our findings in Can 10 cells contrast this view, likely because Can 10 cells, like developing hepatocytes, form tubular BCs requiring planar divisions, whereas WIF-9B and HepG2 hepatocytes used previously cannot form tubular BCs and undergo non-planar divisions. Furthermore, we identify ARHGEF17 as the RhoGEF that localizes to AJs in an E-cadherin- and F-actin-dependent manner, activates RhoA, and is required for BC elongation. Together, these data provide a mechanistic framework for how E-cadherin drives BC elongation by coordinating spindle orientation and RhoA activation.

In contrast, N-cadherin localizes exclusively to linear AJs and inactivates RhoA via the p120- catenin isoform ARVCF and its partner p190B/ARHGAP5. Thus, E-cadherin activates RhoA through ARHGEF17 during division to promote BC elongation, whereas N-cadherin suppresses RhoA activity post-cytokinesis cells through p190B to prevent sustained ROCK activity, which would otherwise stabilize the actomyosin belt and enforce columnar polarity (Fig. 7 A) (Lazaro-Dieguez and Musch, 2017). In this way, E- and N-cadherin regulate RhoA activity in a spatiotemporally controlled and opposing manner, enabling BC elongation while maintaining hepatic polarity.

How do cells differentially use E- and N-cadherin to regulate RhoA? The p120-catenin family appears central. Both p120-catenin and ARVCF suppress RhoA via interactions with p190A/ARHGAP35 or p190B/ARHGAP5 (Cho et al., 2010; Ponik et al., 2013; Wildenberg et al., 2006). p120-catenin binds cadherin juxtamembrane regions to stabilize them by preventing endocytosis (Wildenberg et al., 2006). Consistent with this, p120-catenin knockdown in Can 10 cells destabilizes E-cadherin, reducing its level by nearly half. Since E-cadherin promotes RhoA activation via ARHGEF17, p120-catenin-dependent stabilization likely enhances this pathway, further amplified by ROCKs, which bind to p120-catenin (Smith et al., 2012; Smith et al., 2011). In contrast, p120-catenin depletion did not markedly affect N-cadherin levels, consistent with their distinct localization and largely homo-oligomeric mechanism of action in Can 10 cells. Although p120-catenin may contribute to RhoA inactivation, its dominant function appears to be in the E-cadherin pathway, as its knockdown phenocopies E-cadherin depletion. ARVCF, by contrast, specifically associates with N-cadherin. ARVCF knockdown did not alter either E- or N-cadherin levels, but induced RhoA activation and promoted columnar polarity, underscoring its role in mediating RhoA suppression during hepatic polarity establishment and maintenance. Taken together, these findings indicate that through distinct p120-catenin family members, E- and N-cadherin exert opposing effects on RhoA activity, coordinately regulating BC elongation and hepatic polarity.

### Distinct roles of E- and N-cadherin in the formation of nascent cell-cell contacts at the cell division site

The establishment of nascent cell-cell contacts between daughter cells immediately after cytokinesis remains incompletely understood (Osswald and Morais-de-Sa, 2019). In Can 10 cells, this process is primarily mediated by E-cadherin, the only cadherin observed on the entire PM during cell division, including AJs and the cleavage furrow, whereas N-cadherin predominantly accumulates in linear AJs. Because the emergence of hepatic polarity and primordial BC formation are spatially linked to cytokinesis (Wang et al., 2014), these distinct localization patterns support the preferential roles of E-cadherin in polarity establishment and N-cadherin in polarity maintenance.

The differential distribution of E- and N-cadherin may reflect their associations with distinct p120-catenin isoforms. Specifically, p120-catenin and ARVCF exhibit localization patterns similar to E- and N-cadherin, respectively. Unlike ARVCF, p120-catenin lacks a typical C-terminal PDZ ligand, a feature of the p120-catenin family lineage (Carnahan et al., 2010). In contrast, ARVCF still retains this ligand motif, which binds to ZO-1 and ZO-2 (Carnahan et al., 2010; Kausalya et al., 2004), likely restricting N-cadherin to regions near tight junctions (Fig. S6 D). As ZO-1 localizes to the division site during the terminal phase of cytokinesis (Wang et al., 2014), ZO-ARVCF-N-cadherin interactions may explain why hepatocyte polarization and primordial BC formation can still occur when E-cadherin is depleted.

We also discovered that the localization of E-cadherin to nascent cell-cell contact sites depends on the accumulation of RhoA–ROCK at the cleavage furrow. Attenuation of RhoA at this region through NuMA or ARHGEF17 knockdown or ROCK inhibition impaired E-cadherin accumulation before cell-cell contact formation. This finding aligns with previous studies indicating that actomyosin contributes to cadherin clustering in an adhesion-independent manner (Campas et al., 2024; Wu et al., 2015). Since ROCK1 serves as a scaffold linking RhoA to p120-catenin (Smith et al., 2012), which is crucial for the constitutive cis-dimerization of E-cadherin at the PM (Vu et al., 2021), it is likely that ROCK and p120-catenin cooperate to facilitate the recruitment of E-cadherin to the nascent cell-cell contacts during cytokinesis. Another potential factor in this process is the basolateral polarity protein Scribble, which has been shown to play a role in E-cadherin recruitment to the daughter cell interface during cytokinesis (Chann et al., 2023). Although the precise mechanism by which Scribble facilitates E-cadherin recruitment remains unclear, its function as a scaffold that stabilizes the E-cadherin–p120-catenin complex likely explains its involvement in this process (Lohia et al., 2012).

### Underappreciated relationships between cytokinesis regulators and BC elongation

This study also revealed that NuMA plays a critical role in BC elongation not only by regulating spindle orientation, but also by promoting E-cadherin recruitment to the daughter cell interface during cytokinesis for nascent AJ assembly. NuMA and E-cadherin exhibit an interdependent relationship: NuMA concentrates at the polar cortex in an E-cadherin-dependent manner, while E-cadherin concentrates at the cleavage furrow in an RhoA-dependent manner. NuMA contributes by restricting RhoA to the cleavage furrow. Although E-cadherin is necessary for NuMA clustering at the polar cortex, it is not required for NuMA’s membrane association, as NuMA remains at the PM after E-cadherin depletion. This association likely depends on LGN and Band 4.1 proteins (Kiyomitsu and Boerner, 2021; Lechler and Mapelli, 2021). LGN directly binds the E-cadherin juxtamembrane region, overlapping with the p120-catenin-binding site (Gloerich et al., 2017; Ishiyama et al., 2010). Although LGN was not identified as a proximal interactor of cadherins in our BioID assay, this may reflect a spatiotemporal regulation, with LGN-E-cadherin interactions enriched at the polar cortex of dividing cells but representing a minor fraction compared to E-cadherin-p120-catenin interactions at the population level. Thus, E-cadherin may still cluster NuMA at the polar cortex through LGN and possibly other interacting proteins such as Band 4.1 (Yang et al., 2009). Scribble, reported to stabilize E-cadherin at mitotic cell cortex and required for NuMA recruitment in mammary epithelial cells (Chann et al., 2023), also interacts with E-cadherin and LGN to coordinate polarity with spindle orientation in prostate epithelial cells (Wang et al., 2018). Scribble may therefore act as a hub linking E-cadherin and NuMA in cytokinesis-dependent BC elongation.

NuMA is well established as a spindle assembly and positioning factor (Kiyomitsu and Boerner, 2021). A recent study also implicates NuMA in restricting RhoA to the cleavage furrow in HeLa cells (Sana et al., 2022). Consistently, we found that NuMA is required for concentrating RhoA at the basal side of the cleavage furrow and for BC elongation in Can 10 cells. Conversely, correct activation and localization of RhoA are thought to be important for the polar accumulation of NuMA in HeLa cells (Sana et al., 2022). Supporting this, ARHGEF17 knockdown impaired RhoA activation and caused spindle orientation defects. The mechanism by which NuMA directs RhoA accumulation at the cleavage furrow remains unclear. Given the significant reduction of RhoA at the PM in NuMA-depleted Can 10 cells, NuMA may regulate RhoA activity through specific GEFs or GAPs, which warrants further investigation.

## Materials and methods

### Cell culture

Can 10 and Fao cells were grown in Ham’s F-12K (Kaighn’s) medium (Thermo Fisher Scientific, 21127022) with 5% FBS (Gibco, 16000044) and 0.4% Antibiotic-Antimycotic (Gibco, 15240062). HEK293T cells were cultured in Dulbecco’s modified Eagle’s medium (Thermo Fisher Scientific, 11995065) supplemented with 10% FBS (Gibco, 16000044) and 1% Antibiotic-Antimycotic (Gibco, 15240062).

### Mice

All experimental animal protocols were approved by the Institutional Animal Care and Use Committee of Peking University. C57BL/6 mouse strain was used in this study. All mice were maintained under specific pathogen-free conditions at 23 ± 2 °C with a 12-h day/night cycle. Female mice aged 8 to 12 weeks were mated with adult male mice. The morning the vaginal plug was detected was designated E0.5.

### Mouse liver lobe culture

Mouse liver lobe culture was performed as previously described (Yang et al., 2017) with slight modifications. Liver tissues were cut into 1 mm^3^ cubes. Fetal liver explants were cultured for 2 days in the presence of DMSO (Sigma-Aldrich, D2650) or 1 µM Y27632 (Selleck, S1049).

### Antibodies and Chemicals

Anti-E-cadherin (36/E-cadherin, 610182) and anti-N-cadherin (32/N-cadherin, 610921) monoclonal antibodies were purchased from BD Biosciences; anti-P-Glycoprotein/mdr (C219, MA1-26528) and anti-ARHGAP5 (3L9I3, MA5-38043) mouse monoclonal and anti-ZO-1 (40-2200; 61-7300), anti-ARVCF (PA5-64129), and anti-pan-cadherin (71-7100) rabbit polyclonal antibodies from Thermo Fisher Scientific; anti-N-cadherin (13A9, 14215) and anti-β-catenin (L54E2, 2677) mouse monoclonal and anti-N-cadherin (D4R1H, 13116), anti-catenin δ-1/p120-catenin (D7S2M, 59854), anti-ROCK2 (E5T5P, 47012), and anti-p190-A RhoGAP (C59F7, 2860) rabbit monoclonal antibodies from Cell Signaling Technology; anti- NuMA1 (AD6-1, MABE1807) mouse monoclonal and anti-Partitioning-defective 3/Par3 (07- 330) and anti-LGN (ABT174) rabbit polyclonal antibodies from Millipore; anti-ZO-1 (R40.76, sc-33725) rat monoclonal and anti-p120 (15D2, sc-23872), anti-4.1R (B-11, sc-166759), anti-N-cadherin (13A9, sc-59987), anti-PKCzeta/aPKC (H-1, sc-17781), anti-MDR1 (D-11, sc- 55510), and anti-RhoA (26C4, sc-418) mouse monoclonal antibodies from Santa Cruz Biotechnology; anti-α−tubulin (66031-1-Ig) mouse monoclonal and anti-GFP (50430-2-AP) rabbit polyclonal antibodies from Proteintech; anti-EPB41L2/4.1G (EPR8873(2), ab175928) and anti-Pericentrin (EPR21987, ab220784) rabbit monoclonal antibodies from Abcam; anti-radixin (GTX105408) and anti-E-cadherin (GTX100443) rabbit polyclonal antibodies from GeneTex; Goat anti-E-cadherin (AF748) polyclonal antibodies from R&D Systems; Goat anti- HNF4A (LS-C758303) polyclonal antibodies from LifeSpan BioSciences.

Y27632 (Sigma-Aldrich, Y0503) was used at a concentration of 3 µM for Can 10 cells. For liver explant culture, 1 µM Y27632 (Selleck, S1049) was used. Dithiobis[succinimidyl propionate] (DSP) was purchased from Pierce (22585).

### Plasmid Construction

The cDNAs encoding rat E-cadherin (aa 1–886), N-cadherin (aa 1–906), ARHGEF17 (aa 1–2057), β-actin (aa 1–375), Mzt1 (aa 1–78), Radixin (aa 1–583), and Myl12b (aa 1–172) were obtained by RT-PCR using RNAs prepared from Can 10 cells. The cDNA for TurboID was a gift from Alice Ting (Addgene plasmid # 107173; RRID: Addgene_107173). For expression of TurboID-tagged E-cadherin and N-cadherin, the cDNA encoding TurboID was inserted to the C-terminus of the cadherins with a flexible linker 4×(GGGGS)GGGS. The cDNA for dimericTomato-2×rGBD was a gift from Dorus Gadella (Addgene plasmid # 176098; RRID: Addgene_176098). The obtained cDNAs were inserted into pLJM1 using In-Fusion Snap Assembly (Takara Bio, 638948). All constructs were sequenced to confirm their identity.

### Lentivirus packaging and transduction in Fao and Can 10 cells

Lentivirus was packaged by co-transfection of the pLJM1 plasmids encoding EGFP, E-cadherin–EGFP, E-cadherin–mScarlet, E-cadherin–TurboID, N-cadherin–EGFP, N-cadherin– TurboID, ARHGEF17–EGFP, mScarlet–β-actin, or Myl12b–mScarlet, with the packaging plasmids pMDLg/pRRE, pRSV-Rev, and the VSV-G envelope-expressing vector pMD2.G into HEK-293T cells using Lipofectamine^TM^ 3000 transfection reagent (Thermo Fisher Scientific, L3000015). The medium was changed to DMEM with 10% FBS 12 h after transfection. Lentivirus-containing supernatants were collected at 48 h post transfection. The collected supernatants were filtered through a Millex-HA syringe filter with a pore size of 0.45 µm (Millipore). For transduction of E-cadherin, N-cadherin, Mylb12b, or 2×rGBD in Can 10 cells, the filtered lentivirus-containing medium was added to the cells in Ham’s F-12K medium with 5% FBS. For transduction of E-cadherin–mScarlet in Fao cells or Mzt1–GFP, Radixin– mScarlet, ARHGEF17–EGFP in Can 10 cells, viral supernatants were concentrated using Lenti-X^TM^ Concentrator (Takara Bio, 631231) and added to the cells. Expression of the tagged proteins was assessed 96 h post-transduction in Fao cells and 48 h in Can 10 cells.

### Generation of Can 10 cells stably expressing dimericTomato–2**×**rGBD

Lentivirus packaging and transduction of pLV–dimericTomato–2×rGBD were performed as described above. Puromycin was added to a final concentration of 2.5 µg/mL 48h post-infection for selection. After one week, single clones were isolated by plating TrypLE (Thermo Fisher Scientific)-dissociated cells at 1 cell/100 µL into 96-well plates, each well containing Ham’s F-12K medium with 5% FBS and 2.5 µg/mL puromycin. After 2 weeks of culture, single clones were screened by microscopy for dimericTomato expression.

### Knockdown with siRNA

The pre-designed 25-nucleotide Dicer-Substrate siRNAs targeting rat E-cadherin, N-cadherin, NuMA, LGN, 4.1R, 4.1G, RhoA, ARHGEF17, p120-catenin, ARVCF, p190B/ARHGAP5, and p190A/ARHGAP35 were purchased from Integrated DNA Technologies (IDT). Negative Control DsiRNA (IDT, 51-01-14-04) was used throughout experiments. Can 10 cells plated at 0.9 × 10^4^/cm^2^ were transfected with 4.8 nM siRNA using Lipofectamine^TM^ RNAi MAX Transfection Reagent (Thermo Fisher Scientific, 13778150) and cultured for 72 h in Ham’s F- 12K medium with 5% FBS.

### Immunofluorescence microscopy

Can 10 cells and Fao cells were cultured on 18-mm coverslips (EMS) coated with rat collagen type I (Corning) in Ham’s F-12K medium supplemented with 5% FBS. For Mdr1 staining, cells were permeabilized with acetone for 2 min at 4 °C. For staining of Par3 and NuMA, cells were fixed in 100% methanol for 2 min at 4 °C followed by 1.8% formaldehyde (Santa Cruz Biotechnology, sc-203049A) for 10 min. Fixed cells were washed twice with PBS (135 mM NaCl, 1.3 mM KCl, 3.2 mM Na_2_HPO_4_, 0.5 mM KH_2_PO_4_, pH 7.4), then blocked with 3% bovine serum albumin (BSA) in PBS for 30 min. For RhoA staining, cells were fixed in 10% trichloroacetic acid (TCA) for 20 min at 4 °C. For other antibodies, cells were fixed with 1.8% formaldehyde for 10 min, washed twice with PBS, and permeabilized in PBS containing 0.1% Triton X-100 and 3% BSA. Indirect immunofluorescence analysis was performed using the following primary antibodies: mouse anti-E-cadherin (BD Biosciences, 610182, 1:200), rabbit anti-E-cadherin (GeneTex, GTX100443, 1:200), mouse-anti-N-cadherin (BD Biosciences, 610921, 1:200), mouse anti-N-cadherin (Santa Cruz Biotechnology, sc-59987, 1:200), rabbit anti-N-cadherin (Cell Signaling Technology, 13116, 1:250), rat anti-ZO-1 (Santa Cruz Biotechnology, sc-33725, 1:500), rabbit anti-ZO-1 (Thermo Fisher Scientific, 40-2200, 1:2000), mouse anti-Mdr1 (Thermo Fisher Scientific, MA1-26528, 1:100), rabbit anti-radixin (GeneTex, GTX105408, 1:250), mouse anti-aPKC (Santa Cruz Biotechnology, sc-17781, 1:200), rabbit anti-pericentrin (abcam, ab220784, 1:1000), rabbit anti-Par3 (Millipore, 07-330, 1:1000), mouse anti-NuMA (Millipore, MABE1807, 1:200), rabbit anti-EPB41L2/4.1G (abcam, ab175928, 1:250), mouse anti-RhoA (Santa Cruz Biotechnology, sc-418, 1:200), mouse anti-p120-catenin (Santa Cruz Biotechnology, sc-23872, 1:200), rabbit anti-ARVCF (Thermo Fisher Scientific, PA5-64129, 1:200), rabbit anti-ROCK2 (Cell Signaling Technology, 47012, 1:200). The secondary antibodies and phalloidin used were: Alexa Fluor 405- conjugated goat anti-rat IgG antibodies (abcam, ab175671), Alexa Fluor 488-conjugated chicken anti-rabbit (Thermo Fisher Scientific, A21441) or goat anti-mouse IgG antibodies (Thermo Fisher Scientific, A11001), Alexa Fluor 555-conjugated goat anti-rabbit (Thermo Fisher Scientific, A21428) or goat anti-mouse IgG antibodies (Thermo Fisher Scientific, A21422), and Alexa Fluor 647-conjugated phalloidin (Thermo Fisher Scientific, A22287). Stained samples were washed twice with PBS and mounted with VECTASHIELD PLUS Antifade Mounting Medium (Vector Laboratories) or VECTASHIELD PLUS Antifade Mounting Medium with DAPI (Vector Laboratories). Confocal images were obtained using a spinning disk confocal scanner unit (CSU-X1, Yokogawa) with a Nikon Ti2-U microscope. The microscope was equipped with a CFI Plan Apo 20×/0.75 Lambda objective (Nikon), a CFI Plan 40×/1.30 oil immersion objective (Nikon), a CFI Apo TIRF 100×/1.49 oil immersion objective (Nikon), an ORCA-Quest qCMOS camera (C15550-20UP, Hamamatsu Photonics), and a Stradus Versalase four-wavelength laser system (Vortran Laser Technology). The imaging system was controlled by VisiView (Visitron Systems).

For live cell imaging of Can 10 cells, cells were grown on ibiTreat µ-slide or 35 mm µ- dish (Ibidi) and imaged at 37 °C in a humidified chamber (Okolab; H301-K-FRAME) with 5% CO_2_. Confocal images were acquired every 10 min with 28 z-stacks at a step size of 0.9 µm (Mzt1–GFP/Radixin–mScarlet) or 26 z-stacks at a step size of 0.8 µm (dTomato–2xrGBD) using a spinning disk confocal scanner unit (CSU-X1, Yokogawa) attached to a Nikon Ti2-E microscope equipped with a CFI Plan 40×/1.30 oil immersion objective (Nikon), an ORCA-Quest qCMOS camera (C15550-20UP, Hamamatsu Photonics) or an EMCCD camera (Evolve 512 Delta, Photometrics), and a Stradus Versalase laser system (Vortran Laser Technology).

For imaging of mouse liver tissues and cultured explants, specimens were fixed in 4% paraformaldehyde at 4 °C overnight, then dehydrated and embedded in paraffin. The paraffin embedded-tissues were sliced into 10 µm-thick sections. After rehydration, antigen retrieval was performed by autoclaving the sections in citrate buffer (10 mM citrate, 0.05% Tween-20, pH 6.0) for 10 min. Indirect immunostaining was performed using the following primary antibodies: Goat anti-E-cadherin (R&D Systems, AF748, 1:100), goat anti-HNF4A (LifeSpan BioSciences, LS-C758303, 1:50), mouse anti-MDR1 (Santa Cruz Biotechnology, sc-55510, 1:100), mouse anti-N-cadherin (Cell Signaling Technology, 14215, 1:100), and rabbit anti-ZO-1 (Invitrogen, 61-7300, 1:200). Nuclei were stained with DAPI (Sigma-Aldrich, D9564, 0.5 µg/ml). Images were captured with TCS SP8 confocal microscope (Leica Microsystems) using a Plan-Apochromat 63×/0.75 NA objective lens (Leica Microsystems).

### Super-resolution imaging

Instant Structured illumination microscopy (iSIM) images were acquired using a microscope (model Olympus IX71 inverted microscope, Evident Scientific) equipped with a UPlanSAPO 60×/NA 1.2 water immersion objective (Evident Scientific), an ORCA-Quest qCMOS camera (model C15550-20UP, Hamamatsu Photonics), and a confocal scan head: VisiTech VT-iSIM (VisiTech International, Inc., UK). The imaging system was controlled by MetaMorph (Molecular Devices). Obtained images were further deconvolved using Microvolution deconvolution plugin (Microvolution) in ImageJ.

Stimulated emission depletion (STED) microscopy images were captured with TCS SP8 STED 3X (Leica Microsystems) using a Plan-Apochromat 100×/1.40 NA oil-immersion STED white objective lens. Acquired mages were reconstructed using Huygens STED deconvolution software (Scientific Volume Imaging) according to the manufacturer’s protocol.

### Image processing and analysis

All images were processed and analyzed using Fiji/ImageJ (2.16.0/1.54g). To quantify the intensity ratio of E-cadherin or N-cadherin on the polar cortex to adherens junction (AJ), the mean intensity of the polar membrane signal was divided by the mean intensity on AJ. Quantification of the intensity ratio of the cadherins on the basal cleavage furrow or on the basal surface surrounding the midbody to AJ was measured by calculating the ratio of the mean intensity of the indicated basal surface area divided by the mean intensity on AJ. For quantification of the NuMA intensity ratio of polar cortex to the cytoplasm, the mean intensity of the polar membrane signal was divided by the mean intensity in the cytoplasm area between a spindle pole and the polar cortex. Quantification of polar enrichment of NuMA was measured by calculating the ratio of the mean intensity of polar membrane signal divided by the mean intensity of equatorial membrane signal. To quantify the intensity ratio of BC-associated RhoA or active RhoA vs. cytoplasm, the mean intensity of apical membrane signal was divided by the mean intensity in the cytoplasm. Quantification of apical ROCK2 signal was measured by calculating the ratio of the mean intensity of the apical edge delineated by aPKC signal divided by the mean intensity in the cytoplasm. All data analyzed for intensity measurements were from at least two independent experiments. For quantification of ZO-1 length, the length of ZO-1 positive structure was measured. For measurement of spindle angles, the acute angle between the spindle axis and the line along the apical surface was measured. Quantification of cells with BCs was performed as previously described (Wang et al., 2014), with minor modifications. Briefly, cells exhibiting a radixin- or aPKC-positive apical membrane positioned between adjacent cells was classified as having hepatic polarity. BC structures were categorized as follows: a BC formed between two cells was defined as a primordial BC (pBC), while a BC enclosed by three or more cells and exhibiting a long-to-short axis ratio >2.0 was defined as a tubular BC (tBC). Cells with apical domains facing the culture surface were classified as having columnar polarity. The sum of cells with hepatic or columnar polarity was counted as apical-basal polarized cells. All quantifications of BC structures were based on at least four independent experiments.

### Reversible Cross-Link immunoprecipitation (ReCLIP)

The ReCLIP assay was performed as described by (Smith et al., 2011), with minor modifications optimized for Can 10 cells. 2×10^6^ cells were cultured on a 10 cm dish for 72 h in Ham’s F-12K medium supplemented with 5% FBS. Cells were washed twice with Dulbecco’s phosphate-buffered salt solution containing Ca²⁺ and Mg²⁺ (DPBS; Corning, 21030CM) at room temperature (RT). After removal of DPBS, 10 mL of a 0.5 mM Dithiobis[succinimidyl propionate] (DSP) crosslinker (Pierce, 22585) solution in DPBS was added to each plate. Cells were incubated with the crosslinker for 30 min at RT with occasional agitation. Following addition of 10 mL quenching solution (20 mM Tris-Cl, pH 7.6), cells were incubated at RT for additional 10 min. After quenching, cells were washed once with chilled DPBS, and lysed with RIPA buffer (150 mM NaCl, 1 mM Na_2_EDTA, 1 mM EGTA, 1% NP-40, 1% sodium deoxycholate, 2.5 mM Na_4_P_2_O_7_, 1 mM β-glycerophosphate, 1 mM Na_3_VO_4_, 1 µg/ml leupeptin, 20 mM Tris-Cl, pH 7.5; Cell Signaling Technology, 9806) containing protease inhibitor cocktail (Sigma-Aldrich, P8340). Lysates were homogenized by pipetting 10 times and cleared by centrifugation. The cell lysate was incubated with protein G Sepharose (Cytiva, 17061801) conjugated to an anti-GFP antibody (Proteintech, 50430-2-AP), a Negative Control Mouse IgG (Agilent, X0931), an anti-E-cadherin antibody (BD Biosciences, 610182), or an anti-N-cadherin antibody (Santa Cruz Biotechnology, sc-59987) for 2 h at 4 °C with end-over-end rotation. After the beads were washed three times with RIPA buffer supplemented with protease inhibitors, the precipitants were eluted with 2× Laemmli Sample Buffer (Bio-Rad, 1610747) containing the reducing agents 20% 2-mercaptoethanol and 50 mM DTT.

### Proximity labeling assay

The proximity labeling assay was performed as described by (Cho et al., 2020), with modifications optimized for cadherin expression in Can 10 cells. To generate lentiviruses, HEK293T cells were cultured in 10 cm dishes and transfected with 3.2 µg of the lentiviral vector containing the gene of interest and the lentiviral packaging plasmids pMDLg/pRRE (0.8 µg), pRSV-Rev (0.8 µg), and the VSV-G envelope-expressing vector pMD2.G (2.5 µg) using Lipofectamine 3000 transfection reagent (Thermo Fisher Scientific). After 48 h, the cell medium containing the lentivirus was harvested and filtered through a 0.45-μm filter. 3×10^6^ Can 10 cells were infected with the crude lentivirus and cultured in biotin-free medium (Dulbecco’s modified Eagle’s medium supplemented with 10% FBS). For biotin labeling of transduced cells, biotin was added 2 days post-infection. Biotin (100 mM stock in DMSO) was diluted in the biotin-free medium and added to the cells to a final concentration of 50 µM, followed by incubation at 37 °C for 16 h. Labeling was stopped by gentle washing 10 times with chilled Dulbecco’s phosphate-buffered saline containing Ca²⁺ and Mg²⁺. The supernatant was removed, and the pellet was lysed by resuspension in RIPA buffer (150 mM NaCl, 1 mM Na_2_EDTA, 1 mM EGTA, 1% NP-40, 1% sodium deoxycholate, 2.5 mM Na_4_P_2_O_7_, 1 mM β-glycerophosphate, 1 mM Na_3_VO_4_, 1 µg/ml leupeptin, 20 mM Tris-Cl, pH 7.5; Cell Signaling Technology, 9806) containing protease inhibitor cocktail (Sigma-Aldrich, P8340) and incubated for 10 min at 4°C. Lysates were clarified by centrifugation at 15,000 *g* for 20 min at 4 °C. For enrichment of biotinylated proteins, 400 µg streptavidin-coated magnetic beads (Pierce, PI88816) were washed twice with RIPA buffer and incubated with clarified lysates for 2 h at 4 °C with end-over-end rotation. The beads were then washed twice with 1 ml RIPA buffer, once with 1 ml 1 M KCl, once with 1 ml 0.1 M Na_2_CO_3_, once with 1 ml 2 M urea in 10 mM Tris-Cl (pH 8.0), and twice with 1 ml RIPA buffer. After enrichment, biotinylated proteins were eluted by boiling the beads in 40 µl 2× Laemmli Sample Buffer (Bio-Rad, 1610747) supplemented with 20 mM DTT and 2 mM biotin. The eluted proteins were separated by SDS-PAGE gel for further processing and preparation for LC-MS/MS analysis.

### LC-MS/MS analysis and data processing

Eluted proteins were separated on an SDS-gel for approximately 5 mm, fixed, and stained using Colloidal Blue Staining kit (Thermo Fisher Scientific, LC6025). The entire region of the gel-containing protein was excised, reduced with TCEP, alkylated with iodoacetamide, and digested with trypsin. The resulting tryptic digests were analyzed using a Q Exactive Plus mass spectrometer (Thermo Fisher Scientific) coupled with Vanquish UHPLC system (Thermo Fisher Scientific). Mass spectrometry data were searched with full tryptic specificity against the UniProt Rat proteome database (07/21/2022) and a contaminant database using MaxQuant 2.4.2.0. Variable modifications searched include: Acetylation on protein N-terminus, Oxidation on methionine and Deamidation on asparagine. Protein quantification was performed using razor + unique peptides. Common contaminants, incorrect identifications, and proteins identified by only a single razor + unique peptide were removed from the protein list. Protein and peptide abundance was measured using iBAQ (intensity-based absolute quantification), followed by log₂ transformation for normalization prior to limma analysis. Differential analysis was performed using the R package limma (v3.64.1) with the parameter "trend=TRUE". Significant protein identifications between E-cadherin and N-cadherin samples were defined as those showing: (1) minimum fold change (experimental/control) > 2, and (2) adjusted p-value < 0.05. Protein accession numbers were converted to gene symbols using UniProt for subsequent gene ontology enrichment analysis.

### Immunoblot analysis

Can 10 cells and Fao cells were washed with ice-cold DPBS containing Ca²⁺ and Mg²⁺ (Corning, 21030CM) and lysed in 2× Laemmli Sample Buffer (Bio-Rad, 1610747) containing 1× protease inhibitor cocktail (Sigma-Aldrich, P8340) and 10% 2-mercaptoethanol. Western blotting was performed following standard protocols, using the following primary antibodies: Anti-E-cadherin (BD Biosciences, 610182, 1:500), anti-N-cadherin (Cell Signaling Technology, 13116, 1:200), anti-pan-cadherin (Thermo Fisher Scientific, 71-7100, 1:1000), anti-catenin δ-1/p120-catenin (Cell Signaling Technology, 59854, 1:1000), anti-RhoA (Santa Cruz Biotechnology, sc-418, 1:250), anti-ARVCF (Thermo Fisher Scientific, PA5-64129, 1:250), anti-GFP (Proteintech, 50430-2-AP, 1:500), anti-NuMA1 (Millipore, MABE1807, 1:400), anti-LGN (Millipore, ABT174, 1:1500), anti-4.1R (Santa Cruz Biotechnology, sc- 166759, 1:200), Anti-EPB41L2/4.1G (abcam, ab175928, 1:1000), anti-p190-A RhoGAP (Cell Signaling Technology, 2860, 1:500), anti-ARHGAP5/p190B (Thermo Fisher Scientific, MA5- 38043, 1:500), and anti-α-tubulin (Proteintech, 66031-1-Ig, 1:2000). For quantification of E-cadherin and N-cadherin intensity, the membranes were incubated with IRDye 800CW goat anti-mouse (LI-COR Biosciences, 926-32210) and IRDye 680RD goat anti-rabbit secondary antibodies (LI-COR Biosciences, 926-68071) diluted at 1:2000 in Antibody Diluent (Thermo Fisher Scientific) at RT for 1 h. After three washes with 1× Wash Buffer (Thermo Fisher Scientific), the blots were scanned with ODYSSEY M (LI-COR Biosciences). For other antibodies, Rabbit and Mouse Optimized HRP Reagent (Thermo Fisher Scientific) were used as secondary antibodies. The blots were developed with SuperSignal West Femto Maximum Sensitivity Substrate (Thermo Fisher Scientific, 34096) after three washes with the wash buffer, followed by scanning with LI-COR ODYSSEY M. The obtained images were analyzed with Image Studio (LI-COR Biosciences).

For isolation of mouse hepatoblasts and hepatocytes, embryonic mouse liver tissues were digested with a mixture of collagenases (2.5 mg/ml collagenase II, 2.5 mg/ml collagenase IV, dissolved in RPMI 1640 medium) at 37 °C. The cells were labeled with an anti-mouse DLK- FITC (MBL International, D187-4, 1:100) antibody, filtered through a 70 µm cell strainer, and sorted using a BD FACSAria Fusion cell sorter (BD Biosciences). Dead cells were excluded using DAPI staining. The targeted mouse cells were lysed in 2× loading buffer [5× loading buffer (NCM Biotech, WB2001) diluted in strong RIPA lysis solution (50 mM Tris (pH 7.4), 150 mM NaCl, 1% Triton X-100, 1% sodium deoxycholate, 0.1% SDS)]. Western blotting was performed following standard protocol using the following primary antibodies: Anti-E-cadherin (BD Biosciences, 610182, 1:1000), anti-N-cadherin (Cell Signaling Technology, 14215, 1:1000), anti-pan-cadherin (Thermo Fisher Scientific, 71-7100, 1:1000), and anti-β-actin (Abclonal, AC026, 1:5000). For quantification of E-cadherin and N-cadherin intensity, the membranes were incubated with IRDye 800CW goat anti-mouse (LI-COR Biosciences, 926-32210) and IRDye 680RD goat anti-rabbit secondary antibodies (LI-COR Biosciences, 926-68021) diluted at 1:5000 at RT for 1 h. After three times washes with PBS containing 0.1% Tween-20, the blots were scanned with ChemiDoc^TM^ MP Imaging System (Bio-Rad). The obtained images were analyzed with Fiji/ImageJ (2.16.0/1.54g).

### Quantification and statistical analysis

All statistical differences were performed using RStudio (ver. 2024.12.0+467). Quantification of polarity types and BC structures between two groups was analyzed by Welch’s t-test, and comparisons among more than two groups were performed using pairwise t-test with the Holm correction. ReCLIP co-precipitated levels of E- or N-cadherin was compared using Welch’s t-test. Expression levels of E- and N-cadherin in p120 and/or ARVCF knockdown cells were analyzed by pairwise t-tests with Holm correction. ZO-1-positive structure lengths in Fao cells expressing mScarlet vs. E-cadherin–mScarlet were compared using the Wilcoxon rank-sum test. Spindle angles or intensity ratio comparisons between two groups were analyzed by Wilcoxon rank-sum test; comparisons among more than two groups used Wilcoxon rank sum-tests with Holm correction.

## Supplemental material

**Fig. S 1**: Expression profiles of E-cadherin, N-cadherin, and p120-catenin in hepatocytes.

**Fig. S 2**: Shared and distinct roles of E- and N-cadherin in polarity development and BC formation in Can 10 cells.

**Fig. S 3**: Roles of E-cadherin-interacting cytokinetic factors in BC elongation.

**Fig. S 4**: Relationship between spindle orientation and BC inheritance pattern.

**Fig. S 5**: Localization and function of ARHGEF17 in relation to E-cadherin.

**Fig. S 6**: Role of N-cadherin in regulating ROCK2 localization at the apical edge and the influence of ECM signaling on hepatic polarity development.

**Table S1**. Plasmids and siRNA sequences used in this study.

## Data availability

The data supporting the findings of this study are included in the paper and its supplemental information and will be available from the primary corresponding author (Erfei Bi via) upon request.

## Acknowledgments

We would like to thank the members of the Bi and Xu labs for stimulating discussions, the Proteomics and Metabolomics Facility at the Wistar Institute for mass spectrometry work and analysis, and Cell and Developmental Biology Microscopy Core for imaging assistance.

This work was supported by the National Institutes of Health grant DK128861 (to E. Bi), the International Research Fund for Subsidy of Kyushu University School of Medicine Alumni (to J. Hayase), and the National Natural Science Foundation of China grant 32470871 (to L. Yang).

## Author contributions

J.H. and E.B. conceptualized the project; J.H., L.Y., C.R.X., and E.B. designed the experiments; J.H., L.Y., Y.H.Z., and K.W. conducted experiments and analyses; and J.H., L.Y., C.R.X., and E.B. wrote the manuscript.

## Disclosures

The authors declare no competing financial interests.

## Abbreviations

iSIM: instant structured illumination microscopy
LatA: latrunculin A
DSP: dithiobis[succinimidyl propionate]
ReCLIP: reversible cross-link immunoprecipitation

## Notes

### Competing Interest Statement

The authors have declared no competing interest.

