## Supplemental figures S1-S6 and table S1 for "E- and N-cadherin drive hepatic polarity and lumen elongation via opposing effects on RhoA activity"

#### Supplemental Figure 1

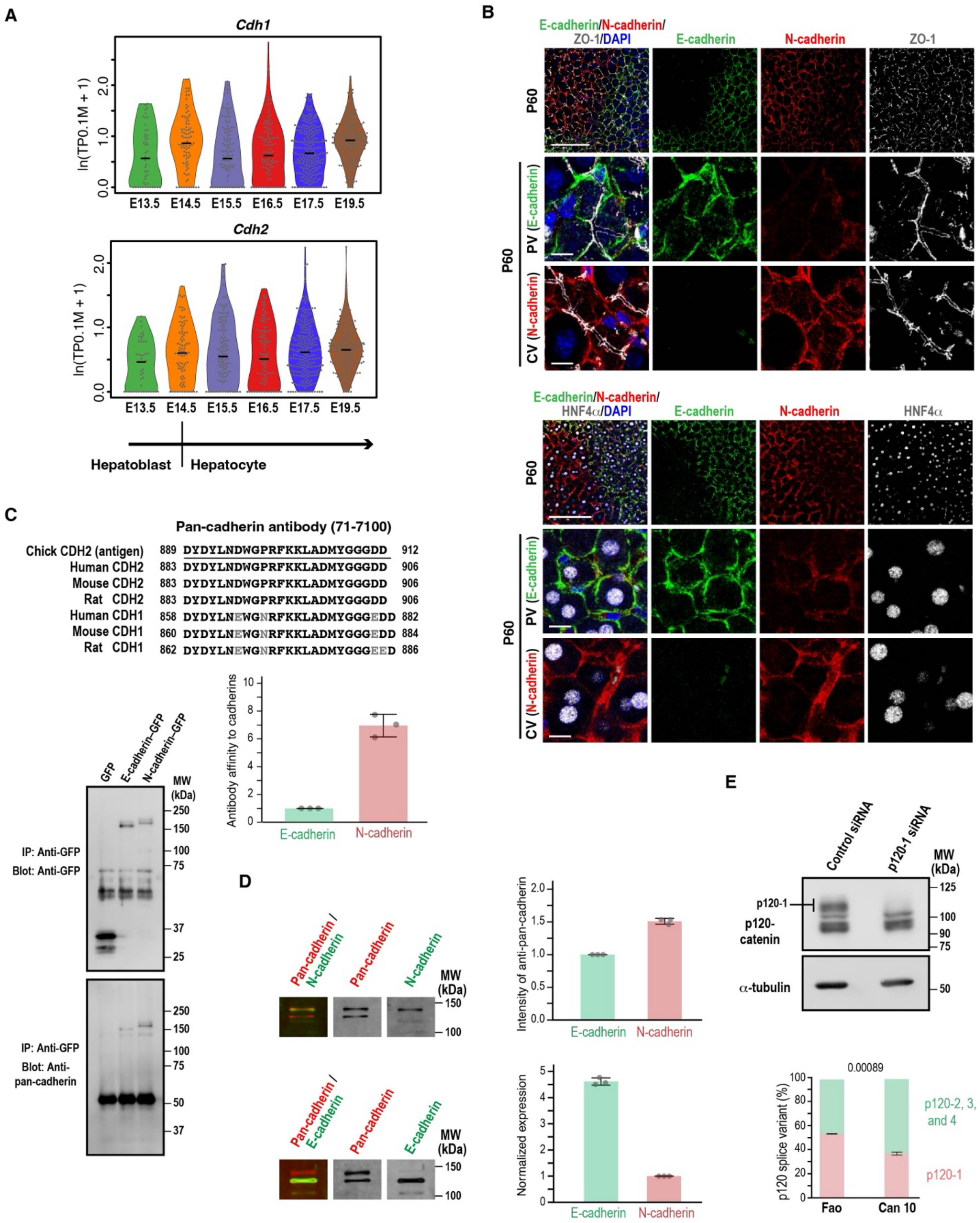

**Figure S1**. **Expression profiles of E-cadherin, N-cadherin, and p120-catenin in hepatocytes**, related to Figure 1.

(A) Single-cell RNA sequencing analysis of cadherin genes during mouse liver development. Violin plots representing the expression of CDH1/E-cadherin and CDH2/N-cadherin in hepatoblasts and hepatocytes. Each dot represents a single cell. The black line within each violin plot indicates the median expression level.

(B) Expression patterns of E- and N-cadherin in adult hepatocytes. Liver sections from P60 were immunostained with antibodies against E-cadherin, N-cadherin, ZO-1, and HNF4α along with DAPI staining. Note the different distributions of E-cadherin and N-cadherin along portal veins (PVs) and central veins (CVs) at P60.

(C) Relative affinities of the anti-pan-cadherin antibody for E- vs. N-cadherin. Left: Lysates from cells expressing GFP-tagged E- or N-cadherin were immunoprecipitated using anti-GFP antibody and analyzed by immunoblotting using anti-GFP and anti-pan-cadherin antibodies. Right: Top—Antigen information for the anti-pan-cadherin antibody (71-7100) and corresponding regions in E-cadherin/CDH1 and N-cadherin/CDH2. Bottom—pulled-down proteins were normalized to GFP intensity, and the ratio of pan-cadherin signal to GFP intensity was calculated. The E-cadherin ratio is set to 1.0. Data represent means ± SD from three independent experiments.

(D) Normalization of E- and N-cadherin expression in Can 10 cells using the anti-pan-cadherin antibody. Top: Raw expression data; the intensity of E-cadherin is set to 1.0. Bottom: Normalized values calculated using the relative antibody affinities determined in (C). Data represent means ± SD from three independent experiments.

(E) Immunoblot analysis of p120-catenin isoform expression in Can 10 cells. Top: Lysates from cells transfected with control or p120-1 siRNA were analyzed using antibodies against p120-1 and a-tubulin (top). Molecular weights (MWs) of marker proteins are indicated in kDa.

Bottom: Quantification of p120-1 vs. other p120 isoforms in Fao and Can 10 cells. Data represent means ± SD from three independent experiments.

Scale bars, 10 μm for the middle and bottom rows and 100 μm for the top row of each image panel (B)

### Supplemental Figure 2

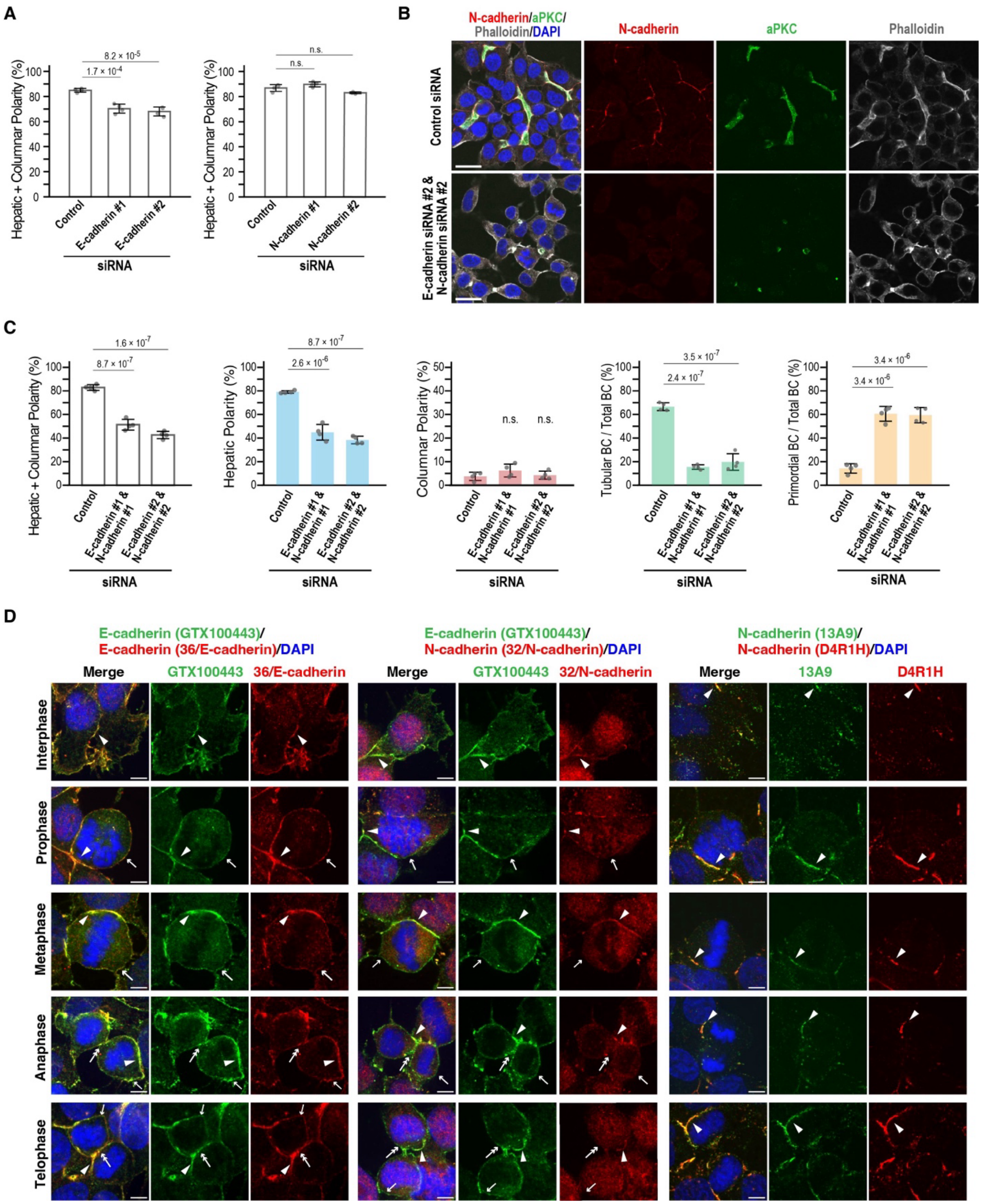

**Figure S2**. **Shared and distinct roles of E- and N-cadherin in polarity development and BC formation in Can 10 cells**, related to Figure 2.

(A) Quantification of total apical-basal polarity (combined hepatic and columnar polarity) in E- or N-cadherin knockdown cells. Data represent means ± SD from four independent experiments (≥480 cells per condition).

(B) E- and N-cadherin jointly contribute to AJ assembly and apical-basal polarity. Representative confocal images of Can 10 cells transfected with E- and N-cadherin siRNAs, cultured for 72 h, and stained with DAPI and antibodies against aPKC and N-cadherin.

(C) Quantification of polarity and BC structures in double knockdown cells. Data represent means ± SD from four independent experiments (≥350 cells per condition).

(D) Localization patterns of E- and N-cadherin during different stages of cytokinesis. Representative confocal images of Can 10 cells cultured for 72 h, and stained with DAPI and the indicated antibodies. Arrowheads, arrows, and double-arrows indicate adherens junctions, polar cortex, and the basal cleavage furrow or midbody, respectively.

Scale bars, 20 μm (B), 5 μm (D).

p-values are indicated at the top of each graph; n.s., not significant.

### Supplemental Figure 3

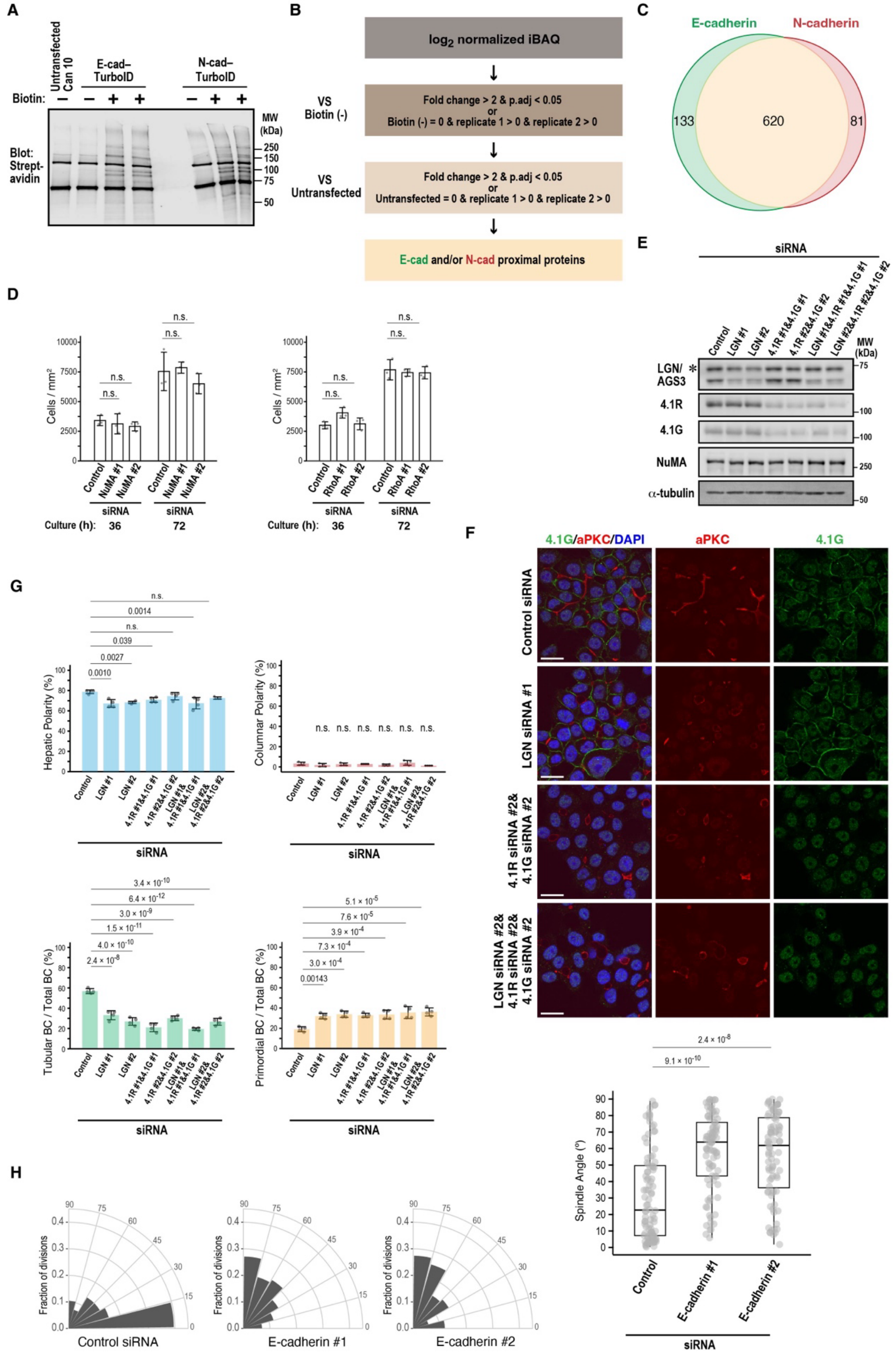

**Figure S3**. **Roles of E-cadherin-interacting cytokinetic factors in BC elongation**, related to Figure 3.

(A) BioID analysis of E- an N-cadherin interactors. Streptavidin blot of Can 10 cells expressing TurboID-tagged E- or N-cadherin. Molecular weights (MWs) of marker proteins are indicated in kDa.

(B) Analysis scheme for the E- and N-cadherin BioID data. Raw intensity-based absolute quantification (iBAQ) values of proteins with ≥2 unique peptides were $\log_2$-normalized and filtered using two sequential criteria. Proteins were retained if, compared to both negative controls [Biotin(-) and Untransfected], they showed (1) significant differential expression [adjusted $p < 0.05$, fold change (FC)>2] or (2) detection in both replicates with complete absence in controls.

(C) Venn diagram of filtered proteins showing overlap between E-cadherin and N-cadherin as well as those specific to each. Numbers in the circles indicate overlapping proteins (620), E-cadherin-specific proteins (133), and N-cadherin-specific proteins (81).

(D) Depletion of NuMA or RhoA does not affect cell proliferation. Can 10 cells transfected with NuMA or RhoA siRNAs were cultured for 36 h or 72 h, stained with DAPI and WGA, and analyzed for cell proliferation. Data represent means ± SD from three independent experiments.

(E) Knockdown efficiencies of LGN, 4.1R, and 4.1G in Can 10 cells. Lysates from siRNA-transfected cells were analyzed by immunoblotting using antibodies against LGN, 4.1R, 4.1G, and α-tubulin. The asterisk marks the AGS3 isoform of LGN. Molecular weights (MWs) of marker proteins are indicated in kDa.

(F) LGN, 4.1R, and 4.1G function in the same pathway to promote BC elongation. Representative confocal images of Can 10 cells transfected with LGN siRNA alone or in combination with both 4.1R and 4.1G siRNAs, cultured for 72 h, and stained with DAPI and antibodies against aPKC and 4.1G.

(G) Quantification of polarity and BC structures in single and double knockdown cells. Data represent means ± SD from four independent experiments (≥742 cells per condition).

(H) E-cadherin depletion alters mitotic spindle orientation. Rose diagrams (left) and combined scatter and box-and-whisker plots (right) show spindle angle distributions relative to the BC (≥84 cells per condition).

Scale bars, 20 µm.

p-values are indicated at the top of each graph; n.s., not significant.

#### Supplemental Figure 4

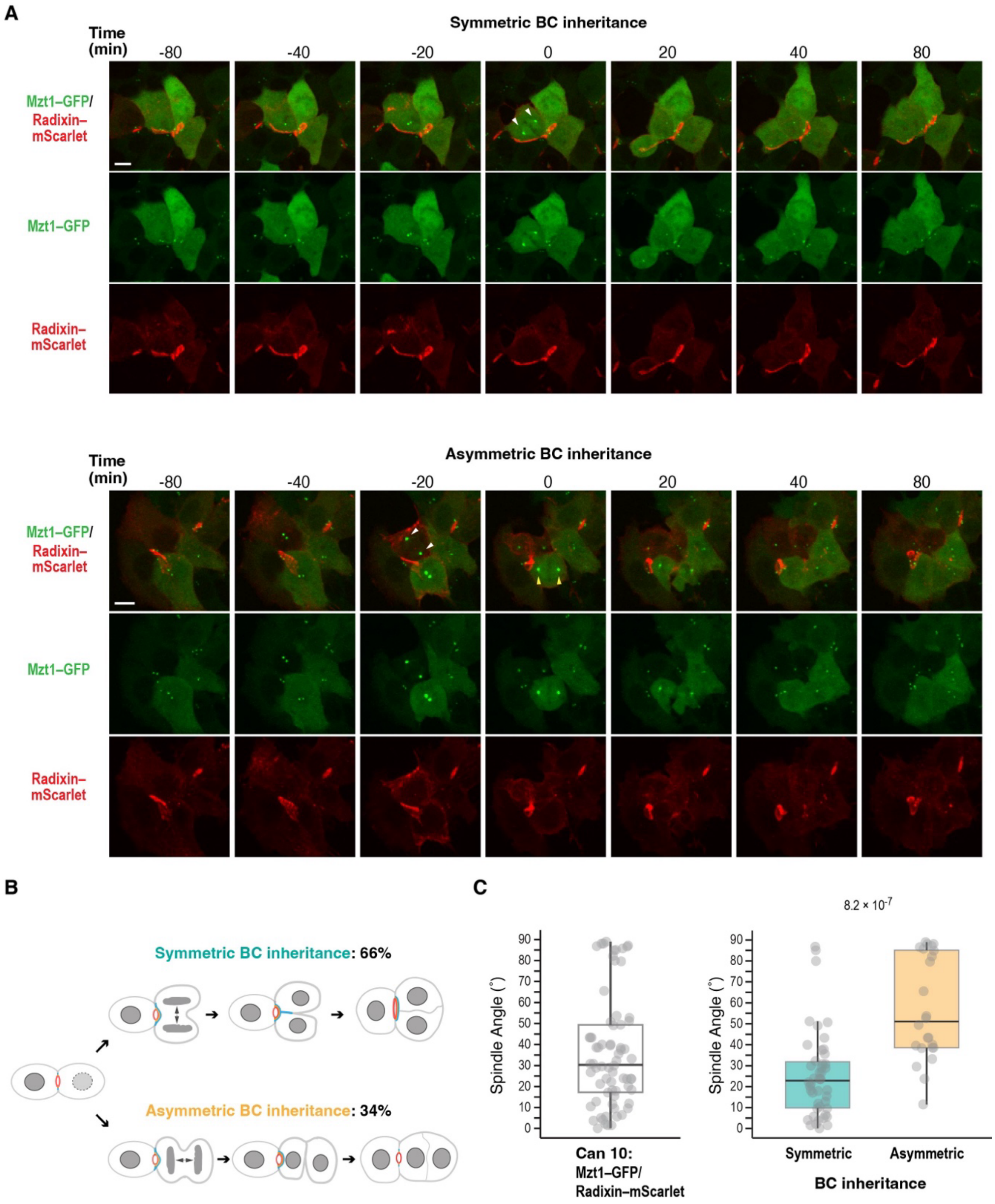

**Figure S4**. **Relationship between spindle orientation and BC inheritance pattern**, related to Figure 3.
**(**A) Montage of time-lapse imaging of Can 10 cells expressing Mzt1–GFP and Radixin–mScarlet during cytokinesis. Maximum intensity projections are shown. The top and bottom panels show symmetric and asymmetric BC inheritance, respectively. White arrowheads at time 0 indicate spindle orientation parallel to the BC, resulting in BC inheritance into both daughters; yellow arrowheads at time 0 indicate spindle orientation perpendicular to the BC, resulting in BC inheritance into only one daughter. Note that Mzt1 brightens during cell division and relocates to the sub-apical region after cytokinesis.
**(**B) Quantification of symmetric versus asymmetric BC inheritance during cell division. Inheritance patterns were assessed by tracking cells expressing Mzt1–GFP and Radixin–mScarlet. Symmetric (emerald green) and asymmetric (gold) inheritance were quantified from 70 cells observed in two independent experiments.
**(**C) Comparison of spindle orientation between symmetric and asymmetric BC inheritance. The combined scatter and box-and-whisker plots on the left show all cells expressing Mzt1-GFP and Radixin-mScarlet; the plots on the right show spindle angle distributions relative to the BC for cells with symmetric or asymmetric BC inheritance (n = 46 cells for symmetric inheritance; n = 24 cells for asymmetric inheritance). Scale bars, 10 µm.
p-value is indicated at the top of the right graph in (C).

#### Supplemental Figure 5

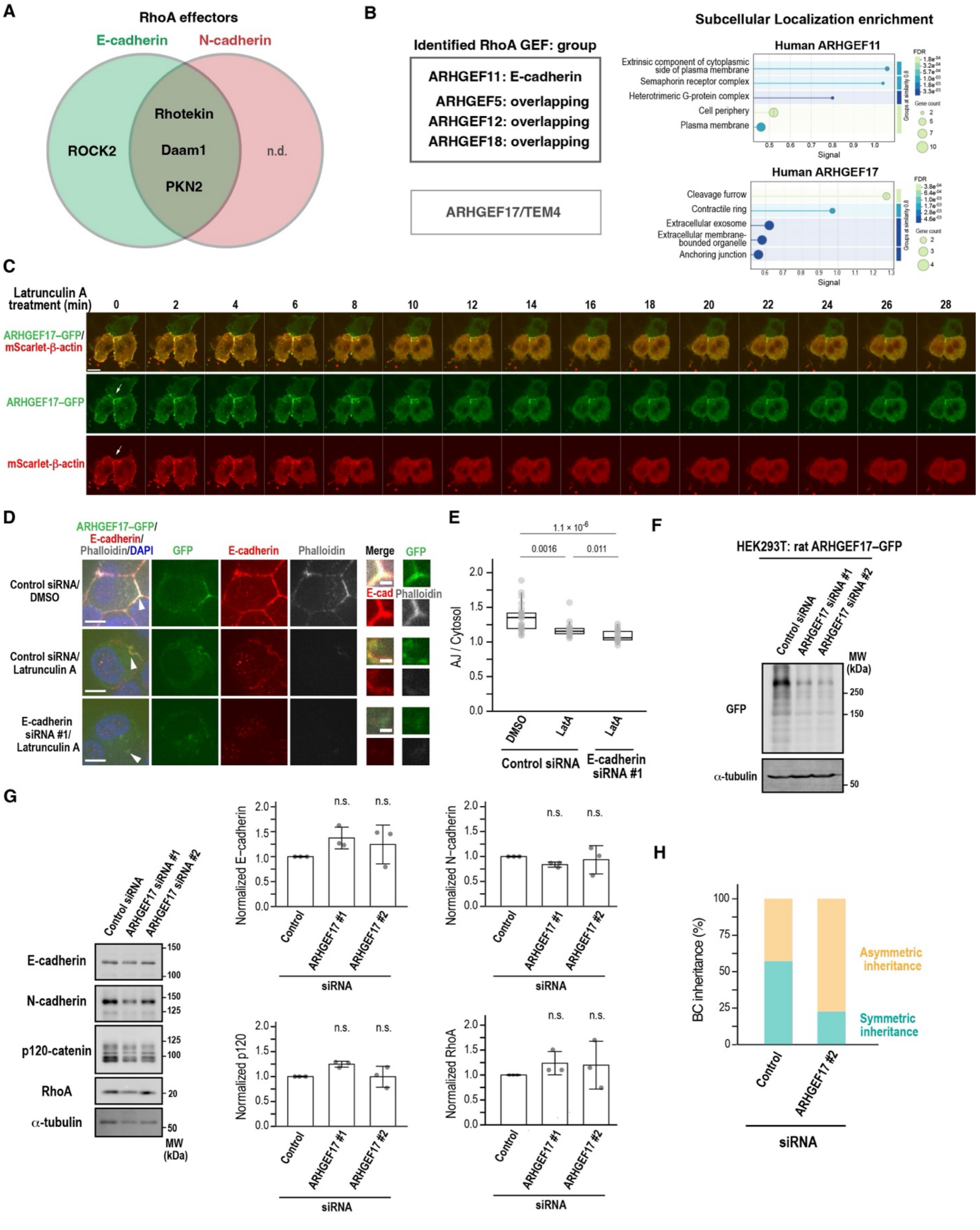

**Figure S5. Localization and function of ARHGEF17 in relation to E-cadherin**, related to Figure 4 and Figure 5.

(A) Venn diagram of RhoA-specific effectors identified by BioID analysis. Note that E-cadherin-specific group contains only ROCK2. n.d., not detected.

(B) RhoA-specific GEFs identified by BioID analysis. ARHGEF17 is shown as an isolated candidate because it was detected in both E-cadherin replicates without a background signal in Biotin (-) but was excluded by the filter due to high background intensity in untransfected cells. GO enrichment analysis of the protein-protein interaction network (STRING) is shown on the right. Note that ARHGEF17 is predicted to localize to the cleavage furrow and contractile ring, whereas ARHGEF11 is not.

(C - E) Localization of ARFGEH17 at AJs depends on F-actin and E-cadherin. (C) Time-lapse montage of Can 10 cells expressing ARHGEF17–GFP and mScarlet–b-actin before and after treatment with LatA. (D) Representative confocal images of Can10 cells expressing ARHGEF17–GFP after treatment with 10 μM LatA for 30 min, combined with control siRNA or E-cadherin siRNA, stained with DAPI, phalloidin, and anti-E-cadherin antibody. The regions indicated by arrowheads are shown as zoomed images. (E) Quantification of ARHGEF17–GFP intensity at AJs relative to the cytoplasm following treatment with Lat A and/or siRNA. Data are from three-independent experiments (≥16 cells per condition).

(F) Validation of ARHGEF17 siRNA efficiency. Immunoblot analysis of ARHGEF17–GFP-expressing HEK293T cells transfected with rat ARHGEF17 siRNAs. Molecular weights (MWs) of marker proteins are indicated in kDa.

(G) ARHGEF17 depletion does not affect protein levels of E-cadherin, N-cadherin, p120-catenin, and RhoA. Left: Representative immunoblots from Can 10 cells transfected with ARHGEF17 siRNAs. Right: Quantification of indicated protein levels normalized to α-tubulin. Data represent means ± SD from three independent experiments; values are normalized to control siRNA-transfected cells.

(H) ARHGEF17 regulates oriented cell division. Proportions of dividing cells expressing dT–2×rGBD that undergo symmetric vs. asymmetric BC inheritance were quantified following transfection with control siRNA or ARHGEF17 siRNA and time-lapse confocal imaging (≥31 cells per condition).

Scale bars, 1 μm (zoomed images in D), 5 μm (D), 20 μm (C).

p-values are indicated the top of each graph; n.s., not significant.

#### Supplemental Figure 6

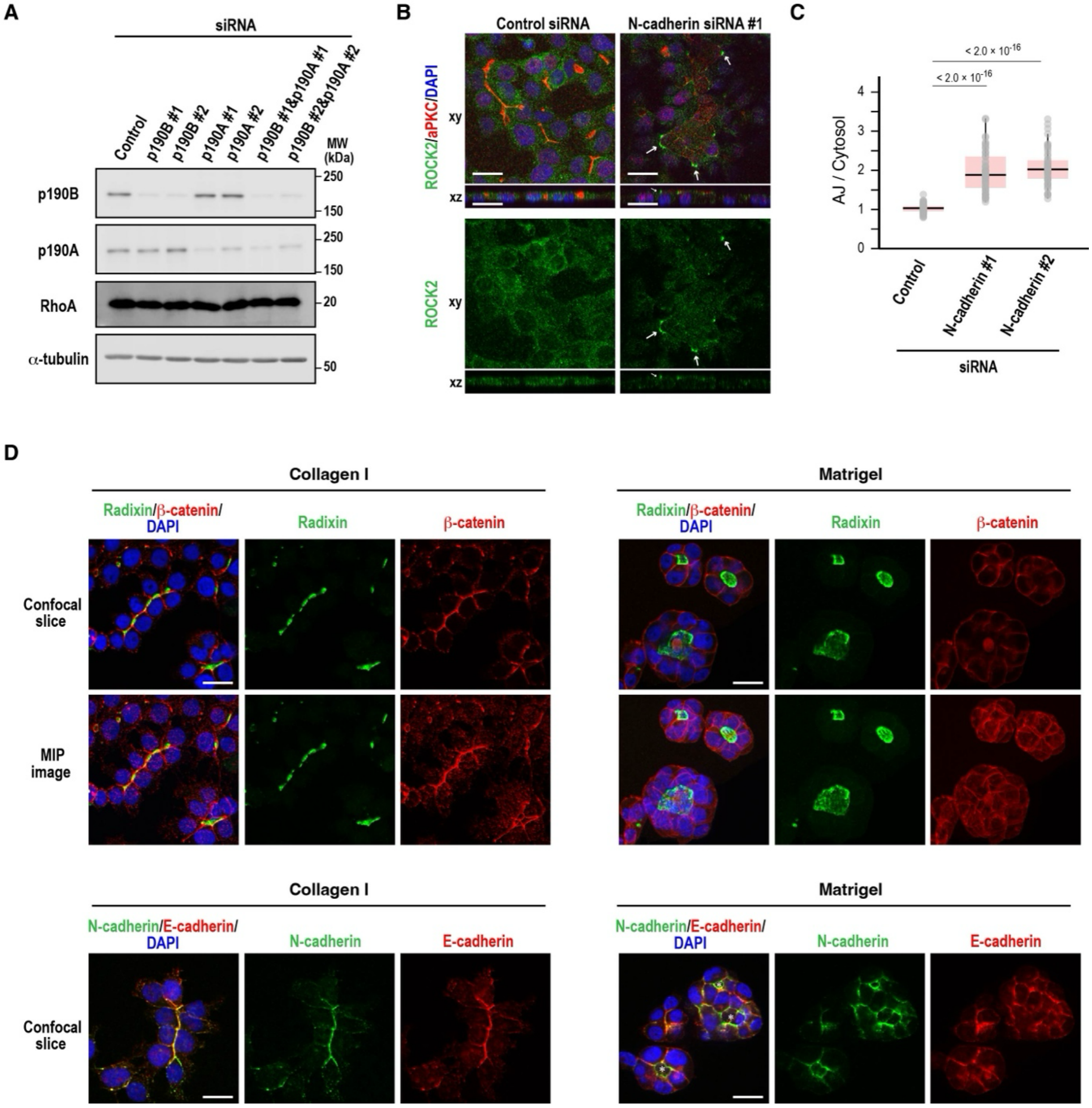

**Figure S6. Role of N-cadherin in regulating ROCK2 localization at the apical edge and the influence of ECM signaling on hepatic polarity development**, related to Figure 6.

(A) Depletion of p190A and/or p190B does not alter RhoA levels. Immunoblot analysis of Can 10 cells transfected with p190A and/or p190B siRNAs. Molecular weights (MWs) of marker proteins are indicated in kDa.

(B - C) N-cadherin depletion increases ROCK2 accumulation at the apical edge. (B) Representative confocal images of Can 10 cells transfected with control or N-cadherin siRNAs, cultured for 72 h, and stained with DAPI and antibodies against aPKC and ROCK2. Arrows indicate ROCK2 accumulation on the apical most junctions. (C) Quantification of ROCK2 intensity at the apical edge vs. cytoplasm. Data are from two-independent experiments (≥54 cells per condition).

(D) Basement membrane components induce cyst-like morphology of Can 10 cells. Representative confocal images of Can 10 cells cultured on collagen type I or Matrigel for 72 h. Cells were fixed and stained with DAPI and antibodies against radixin, b-catenin, N-cadherin, and E-cadherin. Note that N-cadherin prominently localizes to the region surrounding apical lumens (asterisks) in Matrigel-cultured cells, whereas E-cadherin localizes mainly to the basolateral membrane.

Scale bars, 20 μm.

| Oligonucleotide sequences of siRNAs | Target gene |
|---|---|
| E-cadherin #1: GUUGUUGGAAUCUUUUUCUAAAAUG | rat Cdh1 |
| E-cadherin #2: UCACUGUCAAGGAUAUUAAUGACAA | rat Cdh1 |
| N-cadherin #1: GAAUGAAACAACAAGAUUAAAAAAA | rat Cdh2 |
| N-cadherin #2: AGCUCAGCUACACUUGAAUUUUACA | rat Cdh2 |
| NuMA #1: GAGCUGAGGAAUCGGGUCAAGAAUU | rat Numa1 |
| NuMA #2: GAACUGGCCAAGAUGGUAAUGCUAU | rat Numa1 |
| LGN #1: AUUUUACGCUCUCAAGCUAAAAGGA | rat Gpsm2 |
| LGN #2: AAGGCAACAAUCAGUCCGUACUUGA | rat Gpsm2 |
| 4.1R #1: GAGGUGUUCCUUGGAAUUUUACAUU | rat Epb41 |
| 4.1R #2: AGGCUGACUUGGAAUUUCUUGAGAA | rat Epb41 |
| 4.1G #1: GUAUUGAUCUUGGUGACUUCCAGUU | rat Epb41l2 |
| 4.1G #2: GGAGCAUCACACUUUCUACAGACUC | rat Epb41l2 |
| RhoA #1: CAGCCCUGAUAGUUUAGAAAACAUC | rat RhoA |
| RhoA #2: AUGGAGCUUGUGGUAAGACAUGCUU | rat RhoA |
| ARHGEF17 #1: GGACAGUGACGAGCUAAAUUUGGGG | rat Arhgef17 |
| ARHGEF17 #2: CGCUAUGAGCUUCUGGUGAAGGACC | rat Arhgef17 |
| p120-catenin #1: AGAAGGAUUCAGACAGUAAACUUGU | rat Ctnnd1 |
| p120-catenin #2: CAUCUUCCACUGAAGUUUAAUUGUC | rat Ctnnd1 |
| p120-1: GUGGUCCACCUUACUUUUUAUCCUG | rat Ctnnd1 |
| ARVCF #1: GUUUGUGACUUAAAAAUGAAGAAAA | rat Arvcf |
| ARVCF #2: GAGUUAGCUAUUUACUGACAUGGUG | rat Arvcf |
| p190B/ARHGAP5 #1: GAAGCUGACUUGAGAAUUGUCAUGU | rat Arhgap5 |
| p190B/ARHGAP5 #2: GCAACACCAAGUUCUGAUAAAAUGA | rat Arhgap5 |
| p190A/ARHGAP35 #1: GAGAAGCAGAUCAGUCAAGUUCUGA | rat Arhgap35 |
| p190A/ARHGAP35 #2: GAAAUCUGACACUAAGGUACCCUGA | rat Arhgap35 |

| Plasmids | Source |
|---|---|
| pMDLg/pPRE | Addgene #12251 |
| pMD2.G | Addgene #12259 |
| pLJM1-EGFP | Addgene #19319 |
| pRSV-Rev | Addgene #12253 |
| C1(1-29)-TurboID-V5_pCDNA3 | Addgene #107173 |
| dimericTomato-2xrGBD | Addgene #176098 |
| pLJM1–E-cadherin-EGFP | This study |
| pLJM1–E-cadherin–mScarlet | This study |
| pLJM1–E-cadherin–TurboID | This study |
| pLJM1–N-cadherin–EGFP | This study |
| pLJM1–N-cadherin–TurboID | This study |
| pLJM1–ARHGEF17–EGFP | This study |

| | |
|---|---|
| pLJM1–Mzt1–EGFP | This study |
| pLJM1–Radixin–mScarlet | This study |
| pLJM1–mScarlet–β-actin | This study |
| pLJM1–Myl12b–mScarlet | This study |

**Table S1. siRNAs and plasmids used in this study.**
